# ToxoNet: A high confidence map of protein-protein interactions in *Toxoplasma gondii* reveals novel virulence factors implicated in host cell invasion

**DOI:** 10.1101/2021.09.14.460186

**Authors:** Lakshmipuram S. Swapna, Grant C. Stevens, Aline Sardinha da Silva, Lucas Zhongming Hu, Verena Brand, Daniel D. Fusca, Xuejian Xiong, Jon P. Boyle, Michael E. Grigg, Andrew Emili, John Parkinson

## Abstract

The apicomplexan intracellular parasite *Toxoplasma gondii* is a major food borne pathogen with significant impact in children and during pregnancy. The majority of the *T. gondii* proteome remains uncharacterized and the organization of proteins into complexes is unclear. To overcome this knowledge gap, we utilize a biochemical fractionation strategy coupled with mass spectrometry to predict interactions by correlation profiling. Key to this approach is the integration of additional datasets based on gene co-expression as well as phylogenetic profiles that eliminate poorly supported interactions and reduce the number of false positive interactions. In addition to a supervised machine learning strategy, we employed an unsupervised approach in data integration, based on similarity network fusion, to overcome the deficit of high-quality training data in non-model organisms. The resulting high confidence network, we term ToxoNet, comprises 2,063 interactions connecting 652 proteins. Clustering of this network identifies 93 protein complexes, predicting both novel complexes as well as new components for previously known complexes. In particular, we identified clusters enriched in mitochondrial machinery that include previously uncharacterized proteins that likely represent novel adaptations to oxidative phosphorylation. Furthermore, complexes enriched in proteins localized to secretory organelles and the inner membrane complex, predict additional novel components representing novel targets for detailed functional characterization. We present ToxoNet as a publicly available resource with the expectation that it will help drive future hypotheses within the research community.

## BACKGROUND

*Toxoplasma gondii,* the causative agent of toxoplasmosis, is an apicomplexan intracellular parasite of biomedical importance, estimated to infect 30% of the world’s population^1^. It is the leading cause of infectious retinitis in children and is life-threatening in pregnancy and to the immunocompromised^2–4^. Despite its impact, viable vaccines have yet to be developed and few treatments are available. Further, resistance to front line drugs (sulfonamides) is emerging^5^. Able to form tissue-cysts, *Toxoplasma* exhibits a heteroxenous lifestyle, with a sexual phase occurring in the intestinal epithelium of cats and an asexual phase capable of infecting any nucleated cell of any warm-blooded animal. To fulfil its life cycle, *Toxoplasma* is exquisitely adapted to exploit its hosts. For example, to invade its hosts requires specialized processes that mediate host cell attachment, penetration and modulation of host pathways that prevent parasite clearance. Driving this process are hundreds of invasion-related proteins and complexes, that directly impact virulence^6–13^. Key systems include the inner membrane complex (IMC) which drives motility and cell division^14^, Sag-1 related sequence (SRS) proteins which are parasite surface receptors involved in host cell recognition and attachment^13, 15^, microneme proteins, that are also involved in gliding motility and host cell attachment^16^, and dense granule and rhoptry proteins which are secreted during and after host cell invasion to both help the parasite enter the host cell and modulate host cell behaviour subsequently^17–21^. Genome analyses of *Toxoplasma* strains, exhibiting different virulence phenotypes, reveal many of the genes encoding these proteins exhibit significant genetic variation^22^. To better understand the involvement of these genes in pathways driving host cell invasion, a number of approaches such as protein microarrays^23^, CRISPR screens^24^, phosphoproteome analysis^25, 26^, co-expression analysis^6^ and metabolic modeling^27^ have been applied to reveal the contribution of many of these proteins to pathogenesis. However, information on how these proteins are organized into physical protein complexes is lacking. Such an understanding is important as it allows functions to be ascribed to otherwise uncharacterized proteins through *guilt by association*^28^, as well as providing mechanistic insights into how proteins are coordinated to perform specialized biological processes.

Over the past two decades, a number of methods have been developed to help elucidate the physical protein-protein interactions that define protein complexes, including yeast two-hybrid (Y2H) screens^29^, affinity purification mass spectrometry (AP-MS)^30^, spatially restricted enzyme labelling techniques (e.g. BioID^31^ and APEX^32^), phage display^33^ and protein microarrays^34^. For example, in high throughput applications, AP-MS has been applied to generate large scale maps of protein-protein interactions complexes for both yeast and *E. coli*, each comprising hundreds of protein complexes^35, 36^. However such approaches are labor intensive and expensive, requiring the generation of thousands of cell lines carrying tagged gene constructs. Instead, biochemical co-fractionation (or co-elution)^37, 38^, has emerged as a cost-effective route for elucidating protein complexes. In a typical application, proteins are separated into hundreds of biochemical fractions, each subjected to shotgun proteomics. Co-eluting proteins identified in the same fractions are then considered to form stable interactions. Applying this approach has resulted in the recovery of hundreds of stably associated soluble protein complexes for humans^37^ and metazoans^38^. More recently, this strategy has been applied to generate smaller sets of complexes for *Trypanosoma brucei*^39, 40^ and *Plasmodium*^41^. Since co-elution can result in protein pairs that do not physically associate (false positive interactions), computational scoring procedures that integrate additional supporting functional association evidence (e.g. based on expression profiles or literature evidence) are required to increase the overall quality of the predicted interactions.

In this study, we applied a co-elution strategy to generate the first genome-scale physical protein interaction network for *T. gondii*. Based on biochemical cofractionation data, we apply a novel computational strategy, to integrate additional functional data, including coexpression, phylogenetic profiles and domain-domain interactions. Since non-model organisms often lack the depth of known protein complexes of model organisms, required for training supervised machine learning approaches, we complement an approach based on Random Forest classifiers, with an unsupervised approach based on Similarity Network Fusion (SNF)^42^, which provides superior performance in the absence of training data. Together these approaches yield a network of 3,753 interactions capturing 792 proteins. We use this network to define distinct protein complexes including those that have previously been characterized, as well as novel complexes involving invasion proteins that represent novel targets for detailed functional characterization.

## RESULTS

### Co-elution profiling coupled with supervised machine learning recapitulates known protein complexes

This study is the first effort to generate a genome-wide protein-protein interaction network for the apicomplexan parasite *Toxoplasma gondii*, using a biochemical co-elution approach, augmented by the integration of functional genomics datasets. To generate co-elution profiles for *T. gondii* ME49 tachyzoites harvested from human foreskin fibroblasts, we performed six fractionation experiments: five using beads featuring different surface selectivities followed by HPLC^43^, and one using high performance mixed-bed IEX (**Figure 1A; Supplemental Table 1**), resulting in a total of 420 fractions. The proteins in each fraction are cleaved into peptides, which are identified by LC-MS at a false discovery rate of 5%, yielding a total of 1423 unique *T. gondii* proteins (**Supplemental Table 2**). One-half of these proteins are annotated with Gene Ontology (G.O.) terms^44^, while another one-third are defined as ‘hypothetical proteins’ (**Figure 1B**). Of these, 105 represent invasion related proteins, including 17 IMC proteins, 9 rhoptry neck proteins (RONs), 17 microneme proteins (MICs), 27 rhoptry proteins and 35 dense granule proteins (GRAs). Although ∼400 proteins are identified at relatively low abundances (in terms of their spectral counts), ∼1000 proteins possessed ≥ spectral counts (**Figure 1C**). To define interactions between these proteins, the co-elution profiles of the proteins were used to generate six complementary scores for each possible pair in an experiment: Pearson correlation coefficient with noise modeling (PCCNM); weight cross-correlation (WCC); co-apex; mutual information; topological overlap similarity based on PCCNM; and topological overlap similarity based on WCC (**Supplemental Table 3;** see Methods for detailed description). These scoring schemes were used individually for the 6 experiments and once for the elution profiles combining all experiments, which generates a stronger profile for proteins eluting in multiple experiments. To these sets of scores, we integrated additional functional genomics datasets including: two scoring schemes based on gene-co-expression datasets^6^; two scoring schemes based on phylogenetic profile datasets^45^; and a single scoring scheme based on domain-domain interactions^46^ within a supervised machine learning framework (RandomForest – see Methods). For training purposes we used a gold standard set of positive interactions curated from orthologues of known protein complexes collected from the CORUM^47^ and Cyc2008^48^ databases, together with interactions inferred from the Toxocyc resource^49^ and G.O annotations^50^ (**Supplemental Table 4**). Gold standard negative interactions were generated based on differences in cellular localization (see Methods). Integrating datasets within the RandomForest framework gave superior performance relative to any individual dataset (**Figure 1D**). Furthermore, precision-recall curves reveal that the combination of co-elution and functional genomics datasets contributes to the highest recall and precision. **(Supplemental Figure 1).**

**Figure 1.**
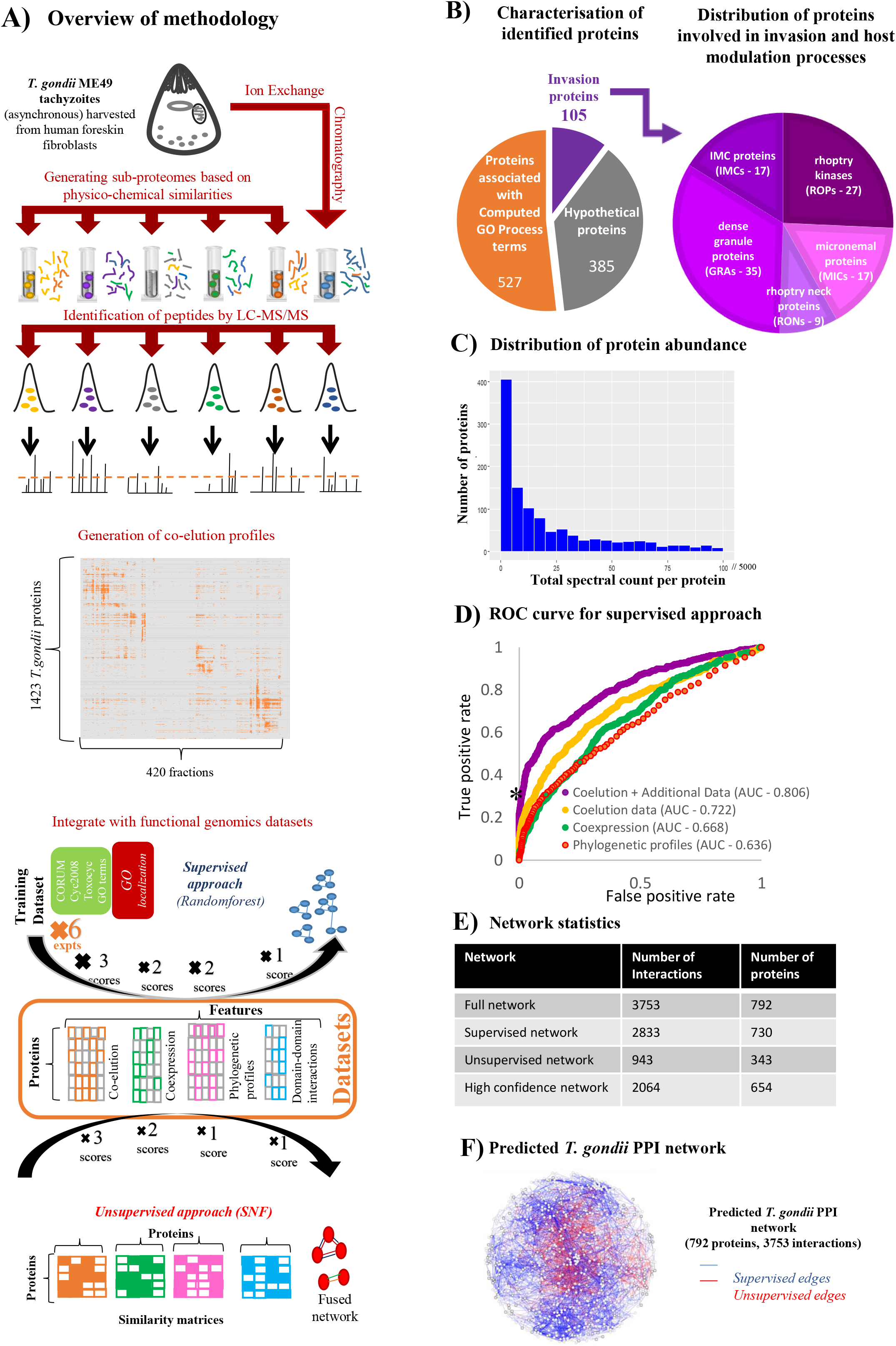
Network generation, statistics and overview analyses. (A) An outline of the methodology employed in this study for generating the predicted *T. gondii* network. (B) Functional characterisation of the proteins identified by co-fractionation in this study, with special reference to invasion clusters. (C) Distribution of protein abundance for co-fractionated proteins. (D) Receiver operating characteristic (ROC) curve for supervised machine learning using RandomForest. The RandomForest output corresponding to a false positive rate of ∼0 and true positive rate of ∼0.3 on the graph (indicated using a * - value of 0.57) was used as the cutoff for selecting the high confidence network. (E) Network statistics of the full network and high confidence network, with a breakup into supervised and unsupervised networks for the full network. (F) The predicted protein-protein interaction network of *T. gondii*.

Based on the set of pairs predicted to be interacting according to the RandomForest approach (corresponding to a score ≥ 5), our approach yielded a network comprising 2,8330 interactions between 730 proteins. From these we used a score cutoff of 0.57 (corresponding to the score at a false positive rate closest to ∼0 on the ROC curve, indicated by a * in Figure 1D) to further define a higher quality dataset comprising 1,541 interactions between 549 proteins (**Supplemental Figure 2; Supplemental Table 5**). This network segregates proteins associated with invasion from known complexes and functional interactions conserved across organisms. As expected, the supervised network recapitulates many known complexes including the proteasome, the ribosome, and a snRNP complex, demonstrating its ability to capture features in the training datasets (**Figure 1C, Supplemental Figure 2**). It also recapitulates several known interactions in *T. gondii*, including MIC1-MIC4-MIC6^51^, ROP5-ROP18^52^, GRA2-GRA4-GRA6^53^.

### Applying an unsupervised machine learning approach identifies additional protein-protein interactions not captured by the supervised approach

A major challenge for inferring protein interactions for non-model organisms is the lack of comprehensive datasets of previously characterized complexes that can serve as training data for machine learning algorithms. To overcome this challenge, we explored an unsupervised machine learning approach, termed Similarity Network Fusion (SNF)^42^, to predict protein interactions in the absence of training data (**Figure 1A**). In this approach, the same scores as used for the supervised approach are used to construct 7 individual networks, based on combinations of three scoring schemes for co-elution data (PCCNM, WCC, Coapex1), two scoring schemes for co-expression datasets (COEXPR-RS, COEXPR-MA), and two scoring schemes for phylogenetic profiles (MI-Pij, MI-PresAbs) (see Methods for more details). Networks are then fused using a method based on message-passing theory to identify interactions supported by multiple datatypes, eliminating poorly supported interactions and strengthening interactions supported by multiple datatypes. We exhaustively explored all possible combinations of parameters to identify the set of protein interactions that captured that had a score greater than the score corresponding to a false positive rate of 0.05 with respect to the training data. From these networks, the network captured the greatest number of likely complexes (corresponding to the highest number of clusters with Bader- Hogue score ≥ .25 (PMID: 22426491), resulting in 943 interactions between 343 proteins (**Supplemental Table 6**) was used. This network was constructed using a combination of the following scores fused by SNF: PCCNM, WCC, Coapex1, Coexpr-RNAseq, Coexpr- Microarray, MI-pij. To further reduce the number of false positives in this dataset, we additionally filtered out interactions with co-expression scores <0.5 (**Supplemental Figure 3**), which corresponds to the cutoff where the supervised network starts losing interactions), resulting in 523 interactions between 282 proteins. This network consists of 73 hypothetical proteins and 32 invasion-related proteins, with some connected interactions localized to the same compartment (eg. MIC4, chitinase-like protein CLP1, TGME49_200270 are localized to the microneme, according to GO evidence codes).

Comparison of the overall supervised and unsupervised networks reveals that 62 proteins are exclusive to the unsupervised network (**Supplemental Figure 4A**). These proteins are involved in 262 interactions, including some paralogs such as HMG_box_containing_protein, aminopeptidases, MIC17A-MIC17B, MIC17A-PAN/Apple domain-containing protein, which are likely to be involved in functional interactions. The distribution of their spectral counts is similar to that of all proteins in the co-elution dataset (**Supplemental Figure 4B**). In terms of the topmost co-elution scores (pcc-noise model, weighted cross correlation, overall coapex score 1), the predicted unsupervised pairwise interactions score significantly better than randomly generated interactions; however, they are worse than that of supervised scores for PCCNM and WCC, but comparable in terms of the coapex score. Of the remaining proteins, 415 are exclusively predicted in the supervised network whereas 315 are common to both (Supplemental Figure 4A). For these common proteins, the two approaches predict 73 common interactions, the rest are mutually exclusive – with 2053 interactions predicted by the supervised approach, and 898 interactions predicted by the unsupervised approach. Based on the six scoring schemes outlined above, we find that the supervised approach captures interactions with higher co-elution scores in general (except for coapex score – which is similar for both supervised and unsupervised approaches) and closer phylogenetic profiles (Pij score) than the unsupervised approach, which still demonstrates improved performance over sets of randomly generated interactions (**Supplemental Figure 4C**). Combining the supervised and unsupervised approaches generates a single *combined* network of 792 proteins and 3,753 interactions (**Figure 1F**). The networks share 315 proteins (an overlap of 43.1% for supervised; 83.1% for unsupervised) and 73 interactions (an overlap of 4.4% for supervised; 8.2% for unsupervised). Among these 792 proteins are 385 annotated as ‘hypothetical’ and 105 predicted to be involved in invasion (**Supplemental Tables 5 and 6**).

From this full network, a high confidence network of 549 proteins and 1541 interactions was derived for the supervised network, based on the score at a false positive rate cutoff of ∼0 on the ROC curve, and a high confidence network of 282 proteins and 523 interactions was derived for the unsupervised network, based on a co-expression score of ≥ 0.5. The combined high confidence version of ToxoNet comprises of predicted interactions from both supervised and unsupervised approaches, resulting in a network of 652 proteins and 2063 interactions, supported by biochemical co-elution (PCCNM (or) WCC scores ≥ 0.5) in at least one of the six experiments, providing a resource of known/putative interactions for several proteins involved in core cellular processes and apicomplexan-specific processes, including 68 invasion-related proteins and 130 hypothetical proteins.

### Integrative analyses validate the biological relevance of ToxoNet

To assess the quality of the network and its ability to recapitulate biologically meaningful relationships, we performed a series of analyses integrating biologically meaningful datasets. Focusing on network statistics, we find that ToxoNet exhibits properties consistent with previously published PPI networks. Notably it exhibits small world properties including a scale-free architecture^54^ and relatively short path length distributions (**Figures 2A and B**). Next we examined the role of essential and conserved proteins in the network. As expected, we find that essential proteins, previously defined through a genome-wide CRISPR screen of the tachyzoite lifecycle^24^, are more highly connected (p = 2.9E-10) and mediate more central roles in the network (p = 1.5E-2) than dispensable proteins (**Figures 2C and D**). These data support the theory that essential proteins serve as network hubs with key organizational roles in PPI networks^55, 56^. Likewise, we find that highly conserved proteins (with orthologs predicted in other, non-apicomplexan, eukaryotes) are also more highly connected than lineage specific proteins, albeit with significant differences only observed with *Toxoplasma*-specific proteins (**Figure 2E**). These findings are consistent with previous studies from many biological networks including *E. coli* and yeast ^35, 57^.

**Figure 2.**
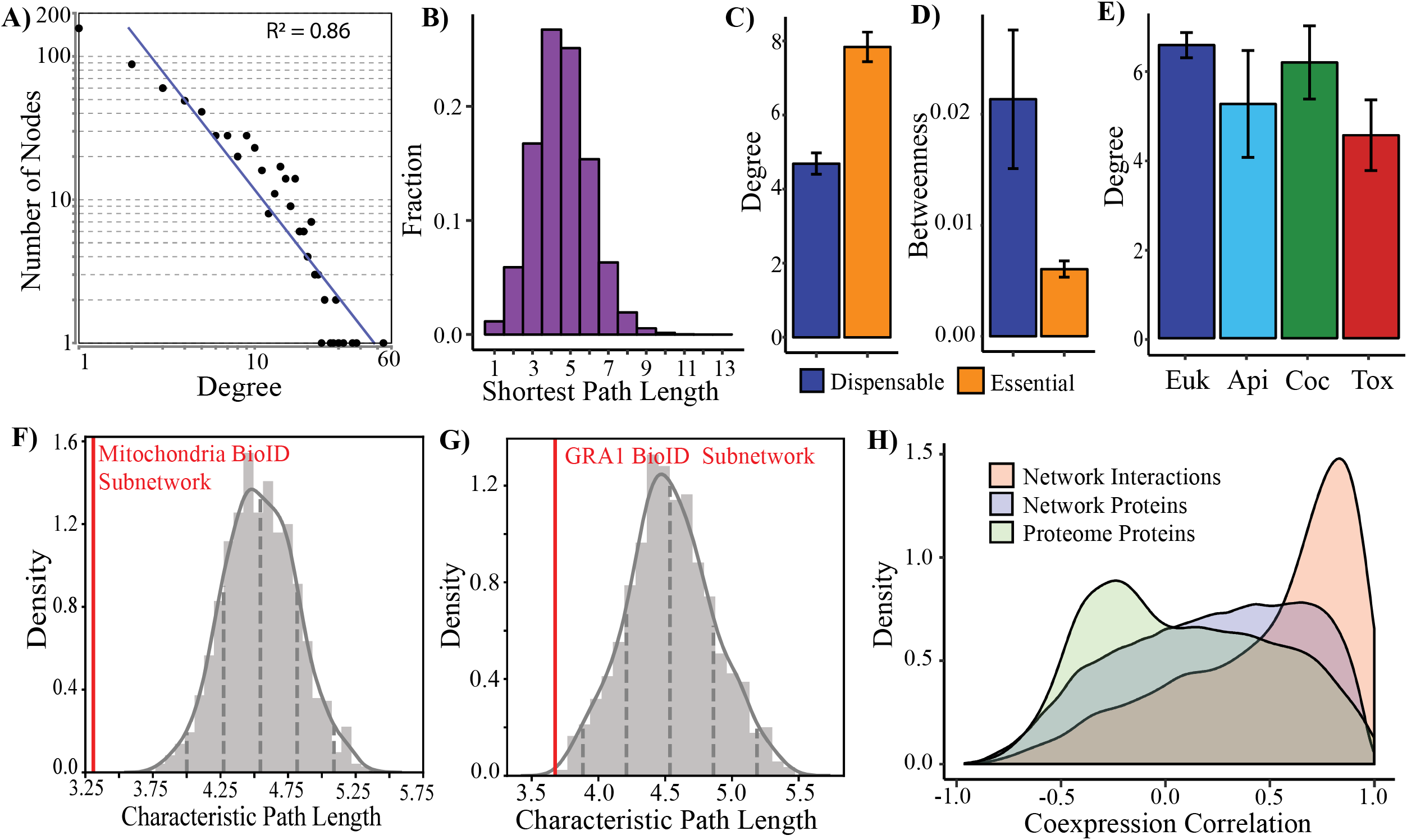
Benchmarking of ToxoNet using previously published datasets. (A) The degree distribution with a power fit line (R^2^ = 0.86). (B) The distribution of shortest path lengths. (C-D) The bar graphs with essential proteins (orange) and dispensable proteins (blue) indicate differences in degree and betweenness centrality (p = 2.9E-10, p = 1.5E-2). (E) The bar graph indicates differences in degree at various evolutionary timepoints (p = 0.02). (F-G) The distribution of random permutations (n=1000) of characteristic path lengths relative to the actual characteristic path length (red line) of a set of candidates identified in BioID experiments as putative mitochondrial (n = 36, F) and dense granule (n = 23,G) proteins ( p= 2.8E-6, p=4.3E-3, respectively). (H) The distribution of coexpression in RNA-seq experiments of interacting network proteins, non-interacting network proteins and a random sampling of the proteome (n=10,000; kolmogorov-smirnov p < 2.2E-16).

We further validated the quality of ToxoNet through comparisons with existing protein interaction data for *T. gondii* as well as other parasites. Recently complementary APEX and BioID approaches^58^ were applied to identify a shared set of 161 *T. gondii* proteins, predicted to localize to the mitochondria. Since we expect co-localized proteins to appear closer in our network, we were reassured by a significantly shorter average pathlength between the 36 proteins that were present in ToxoNet (of the 161 proteins identified in the previous study) than expected by random (average shortest path length = 3.0; p=2.8E-6; **Figure 2F**). A similar dual screen of dense granule proteins^59^ identified a common set of 33 related GRA proteins. Again the average shortest pathlength of the 23 proteins that were found in ToxoNet exhibited a relatively shorter average path length than expected (average shortest pathlength = 3.68; p=4.3E-3; **Figure 2G**). Further datasets based only on a BioID approach^60, 61^, also generally exhibited smaller (but not statistically significant) characteristic pathlengths, with a screen of the IMC-related protein, ISP3, being notable for exhibited a higher average pathlength than random (**Supplemental Figure 5**). This likely reflects the higher false-positive rates associated with these screens in the absence of additional experimental support. The congruency with high confidence datasets demonstrates that the spatial proximity of proteins within the cell is echoed by shorter distance within ToxoNet.

Beyond proximity screens, we also compared ToxoNet with two recently published protein interaction networks for *Trypanosoma brucei*^62^ and *Plasmodium* parasites^41^. Notably both networks were generated using a similar co-elution methodology. Typically protein interaction networks display little overlap between species ^63^. However, here we found that ToxoNet exhibited a significant overlap in protein interactions with both *T. brucei* (283 common interactions, p=6.4E-204) and *Plasmodium* (450 common interactions, p=0; **Supplemental Figure 5B**). While most of these shared interactions occur between highly conserved proteins, and may reflect their inclusion in training data (Supplemental Table 4), we did identify interactions between an apicomplexan-specific protein (TGME49_268830), with an ATP synthase subunit (TGME49_226000) as well as cytochrome c1 (TGME49_246540), to be conserved in the *Plasmodium* network. The former interaction between TGME49_268830 and TGME49_226000 has been validated in *T. gondii*. Its presence in the *Plasmodium* network supports the conservation of this interaction as an apicomplexan-specific adaptation to ATP synthase.

Next we analyzed our network in the context of expression data from RNA-seq datasets that had previously been withheld from the machine learning analyses applied to generate ToxoNet. Pearson correlation values were calculated for protein pairs across eleven time-points encompassing different lifecycle stages, including oocyst^64^, intracellular and extracellular tachyzoites^11^. Expression of putatively interacting proteins has a significantly greater density at higher co-expression correlation values than both non-interacting network proteins (p < 2.2E-16) and random sampling of proteome pair-wise combinations (p < 2.2E- 16; **Figure 2H**). G.O. annotations of putative interactions were also compared using terms annotated to *T. gondii* proteins available on ToxoDB. Despite the limited number of annotated proteins, of those pairs in which both proteins are annotated, 40% have the same

G.O. processes term, 45% have the same G.O. component term and 28% have the same G.O. function term (**Table 1**). It should be noted, however, that G.O. component terms were utilized in the construction of negative training data utilized in supervised learning. Together, these results illustrate that the network properties are consistent with other protein-protein interaction studies.

**Table 1:** Gene Ontology Annotation of Interacting Proteins.

| **G.O.**<br>**Annotation<sup>a</sup>** | **Same Term**<br>**(Network)<sup>b</sup>** | **Same Term**<br>**(Random)<sup>c</sup>** | **p-value<sup>d</sup>** |
| --- | --- | --- | --- |
| Process | 40% | 9% | 4.5E-156 |
| Component | 45% | 11% | 6.5E-86 |
| Function | 28% | 5% | 1.6E-213 |

<sup>a</sup> G.O. annotations were derived from ToxoDB.
<sup>b</sup> The same term (network) column reports the percentage of interactions with both proteins annotated that have the same G.O. term.
<sup>c</sup> The same term (random) column reports the mean percentage of interactions that have the same G.O. term over 1000 random network permutations.
<sup>d</sup> Upper tail z-test.

### ToxoNet recapitulates known protein complexes and identifies novel components and novel complexes

Since co-elution data is enriched for proteins that physically interact, we applied the ClusterONE algorithm^65^ to identify protein complexes based on their interactions (see Methods). In total we identified 93 overlapping clusters (representing putative protein complexes) with an average of 8.2 proteins per cluster (**Figure 3; Supplemental Table 7**). In total, 28 clusters recapitulate 25 training complexes with an overlap score of 0.25 or greater^66^. These include the protein kinase CK2 complex (cluster 72, overlap = 1), the U4/U6.U5 tri-snRNP complex (cluster 33, overlap = 0.79), the C complex spliceosome (cluster 56, overlap = 0.75), the reductive TCA cycle (cluster 54, overlap = 0.75), eIF3 complex (cluster 44, overlap = 0.51), the vacuolar ATP synthase complex (cluster 53, overlap = 0.56) and the box C/D snoRNP complex (cluster 70, overlap = 0.6). Additionally, the ClusterONE algorithm recapitulated experimentally validated complexes absent from training data, including MIC1/4/6 (cluster 83, overlap = 0.34). At a lower cut-off, we also recapitulate GRA2/4/6^11^ (cluster 26, overlap = 0.23), the *T. gondii* ATP Synthase complex with Apicomplexan-specific subunits^67^ (overlapping clusters 12 and 29, overlap = 0.13) and the moving Junction (cluster 88, overlap = 0.13). The glideosome is also partially recapitulated (cluster 52, overlap = 0.11) with GAP45 and MLC1 included in the same cluster as the α β tubulin complex.

**Figure 3.**
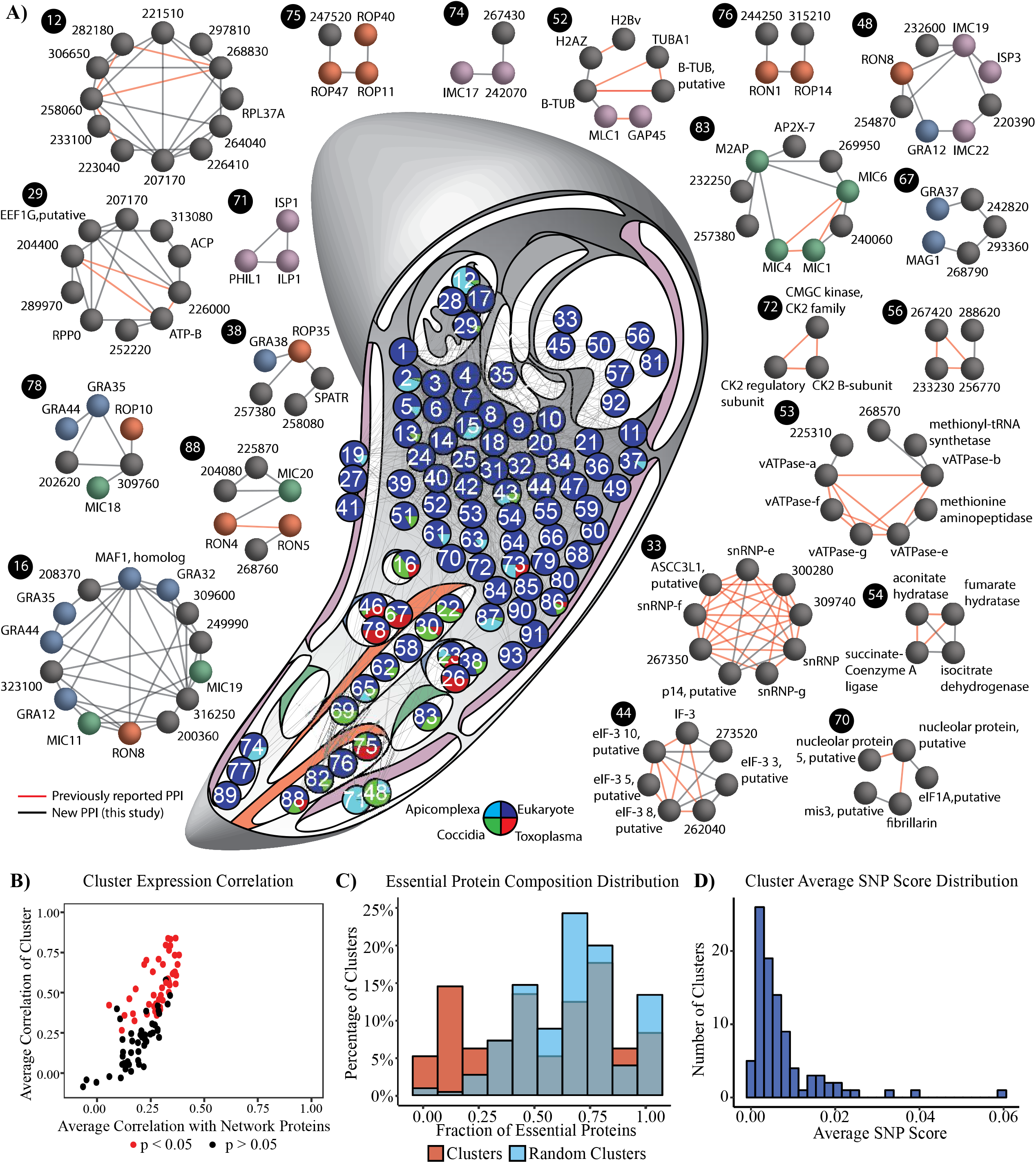
Organization of ToxoNet into discrete protein complexes/. (A) Global organization of predicted complexes is visualized by a graph where nodes represent clusters that are placed in their putative compartment (IMC = purple, micronemes = green; rhoptry = red, dense granules = blue). Nodes are placed in the *T. gondii* schematic in their putative organelle based on their most frequent computed compartment term or known localization. Unclassified clusters are placed in the cytosol. Node coloring displays lineage composition. The edges indicate inter-cluster interactions. Noteworthy clusters are highlighted to the sides with known (red) and novel (black) interactions indicated between proteins. (B) Co-expression of cluster components – clusters are coloured by significance of co-expression. (C) Distribution of essential proteins in clusters and random generated clusters (n=100).

While we found clusters enriched in lineage-specific proteins generally localize to lineage-specific organelles (i.e. rhoptries, micronemes and dense granules), there are also examples of lineage-specific proteins in clusters with proteins that have been shown to be associated with either the cytosol or the mitochondria. Again utilizing expression data withheld from training, we found that of the 93 clusters, 45 (48%) are predicted to be significantly co-expressed **(Figure 3B**). This is consistent with previous studies of protein complexes that display a significant enrichment of component co-expression and likely reflect regulation through common transcription factors^68^. Analysis of essential protein distributions also revealed a non-random pattern of organization (p = 1.079E-06; **Figure 3C**). Specifically, clusters defined by ToxoNet contain a higher proportion of dispensable proteins. Of the 30 clusters enriched in essential proteins (≥ 75% of components are essential), 17 represent recapitulated training complexes. Interestingly, dispensable clusters, composed of ≤ 25% essential proteins (n=25), are significantly enriched in proteins associated with network invasion (i.e., SRS, MIC, ROP, RON and GRA proteins) and IMC proteins (p = 7.3E-35), as well as proteins restricted to Apicomplexa (p = 1.6E-17; **Supplemental Figure 6**). In addition to the high percentage of invasion and IMC proteins per dispensable cluster (41%), these dispensable clusters are composed of an average of 25% hypothetical proteins, offering an exciting wealth of candidates for novel virulence factors. These results are again consistent with previous studies that have shown highly conserved protein complexes to be essential, while those composed of lineage-specific components, reflecting in this case, more recent adaptations to parasitism, tend to be more dispensable.

To further explore the potential impact of these adaptations at the strain level, we leveraged a recent data set of genomes from 80 strains of *T. gondii.* Among proteins identified in our network, we identified 2893 non-synonymous single nucleotide polymorphisms (SNPs) present in at least 4 strains^22, 69^. A SNP score was assigned to each protein based on the number of non-synonymous SNPs normalized by protein length and the average was taken across all cluster proteins to identify putative complexes with the greatest genetic variation (**Figure 3D**). Consistent with previously reported higher rates of SNPs within invasion family proteins, particularly in *GRA* and *ROP* family proteins^22^, we found that putative invasion complexes also exhibited the highest average SNP score (**Supplemental Figure 7**). More interestingly, to predict the SNPs that are most likely to interfere with physical interactions, we examined the Pfam domains [26673716] and identified premature stop codons present in the proteins of the ten complexes with the highest average SNP score. In the experimentally validated MIC1/4/6 complex (cluster 83), we identified five non-synonymous SNPs in three domains mediating interactions with other proteins (G55S and E82Q in the first TSR1 domain of MIC1, N114K andR123L in the EGF2 domain of MIC6 and A171V in the second apple domain of MIC4) ^70, 71^. Notably, strain members of clade E (which includes TgH21016, CASTELLS, TgH26044 and TgH20005) were unique insofar that they carry four of these five mutations, suggesting that the MIC1/4/6 complex in these strains may not occur in its canonical form. Similarly, we found that for 5 strains, including ME49, 3675, B73, PRU, and TgGoatUs21, GRA44 (TGME49_228170) orthologs contain a premature stop codon that results in a 105 amino acid truncation. Given its interactions with other proteins in three predicted invasion-related clusters (16, 23 and 78), we predict these complexes may again exhibit different patterns of organization outside these strains.

### Complex predictions identify novel apicomplexan-specific mitochondrial adaptations

The conserved eukaryotic α-, β-, δ-subunits and the recently identified apicomplexan-specific subunits of the ATP synthase complex^67^ were recapitulated in overlapping clusters 29 and 12, respectively. Strikingly, cluster 12 is enriched in essential apicomplexan-specific proteins, many of which containing predicted mitochondrial targeting sequences (**Figure 4A**). In addition to the four validated ATP synthase subunits (i.e., TGME49_258060, TGME49_268830, TGME49_282180 and TGME49_223040) identified in cluster 12, two additional proteins, that have recently been identified as apicomplexan adaptations to the cytochrome c oxidase (COX) complex: TGME49_264040 and TGME49_297810^58^, were also captured. Recently, TGME49_207170 was identified as a novel component of Complex III of the electron transport chain (ETC)^72^. The remaining two apicomplexan- and coccidian-specific hypothetical proteins, TGME49_306650 and TGME49_221510, remain to be characterized but were identified in both APEX and BioID-based surveys of the *T. gondii* mitochondrial proteome^58^. To assess the localization of TGME49_306650, we engineered a *T. gondii* PRU strain with a 3×HA tag at the C-terminus of the endogenous locus. Mitochondrial localization was validated by colocalization with the mitochondrial marker, F1B ATPase (Figure 4B). Given that this cluster captures two functionally related complexes (i.e. the ATP Synthase and COX complexes), these results indicate that the three remaining uncharacterized apicomplexan specific proteins are likely additional mitochondrial apicomplexan adaptations to oxidative phosphorylation.

**Figure 4.**
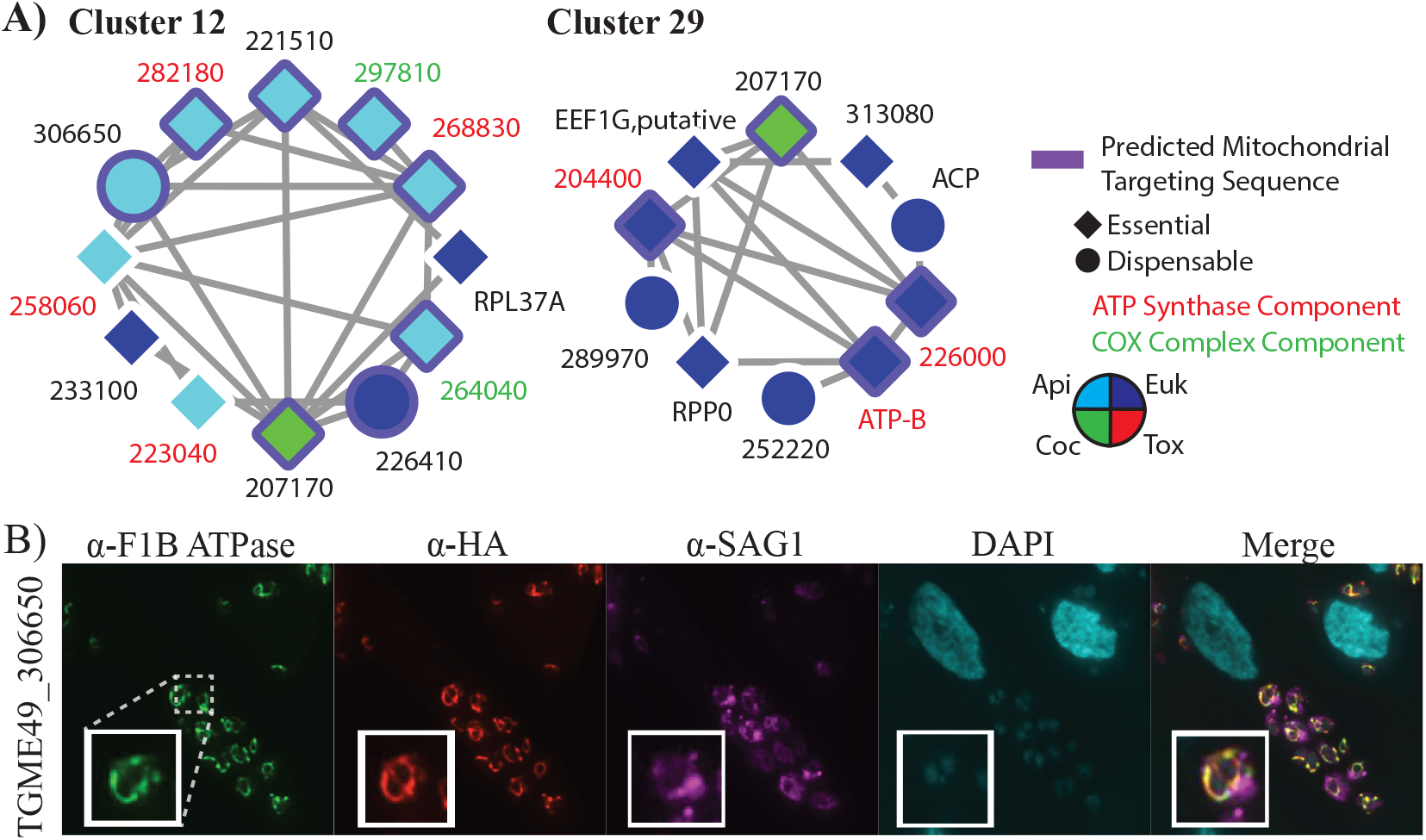
(A) Clusters 12 and Clusters 29 recapitulate mitochondrial complexes: ATP synthase (red text) and the COX complex (green text). The nodes are colored by their lineage and the presence of a purple border indicates a predicted mitochondrial targeting signal by TargetP. Diamond and circle shaped nodes indicate essential and dispensable proteins, respectively. (B) IFA of 306650-3×HA expressing parasites reveals colocalization with the F1B ATPase mitochondrial marker and not the parasite plasma membrane marker SAG1.

### Complex predictions identify novel invasion proteins

ToxoNet predicts 18 clusters representing putative invasion complexes, including 1, 10 and 7 that are predicted to localize to micronemes, rhoptries and dense granules, respectively (e.g. clusters 16, 38, 67, 75, 76, 78, 83 and 86 in **Figure 3A**). Filtering for proteins with validated localization data reveals a set of 38 previously uncharacterized proteins, predicted to be involved in host invasion (**Table 2**). Of these, 15 (39%) are predicted to carry a signal peptide, consistent with their trafficking through the secretory pathway, while 45% are specific to the Apicomplexa lineage. Nine of these proteins have been localized to invasion organelles in ToxoDB user comments, of which two were localized to the IMC. These clusters also predict interactions between well characterized invasion factors and those with little known functional information beyond their localization in discovery screens (e.g., ROP11 and GRA32).

**Table 2:**
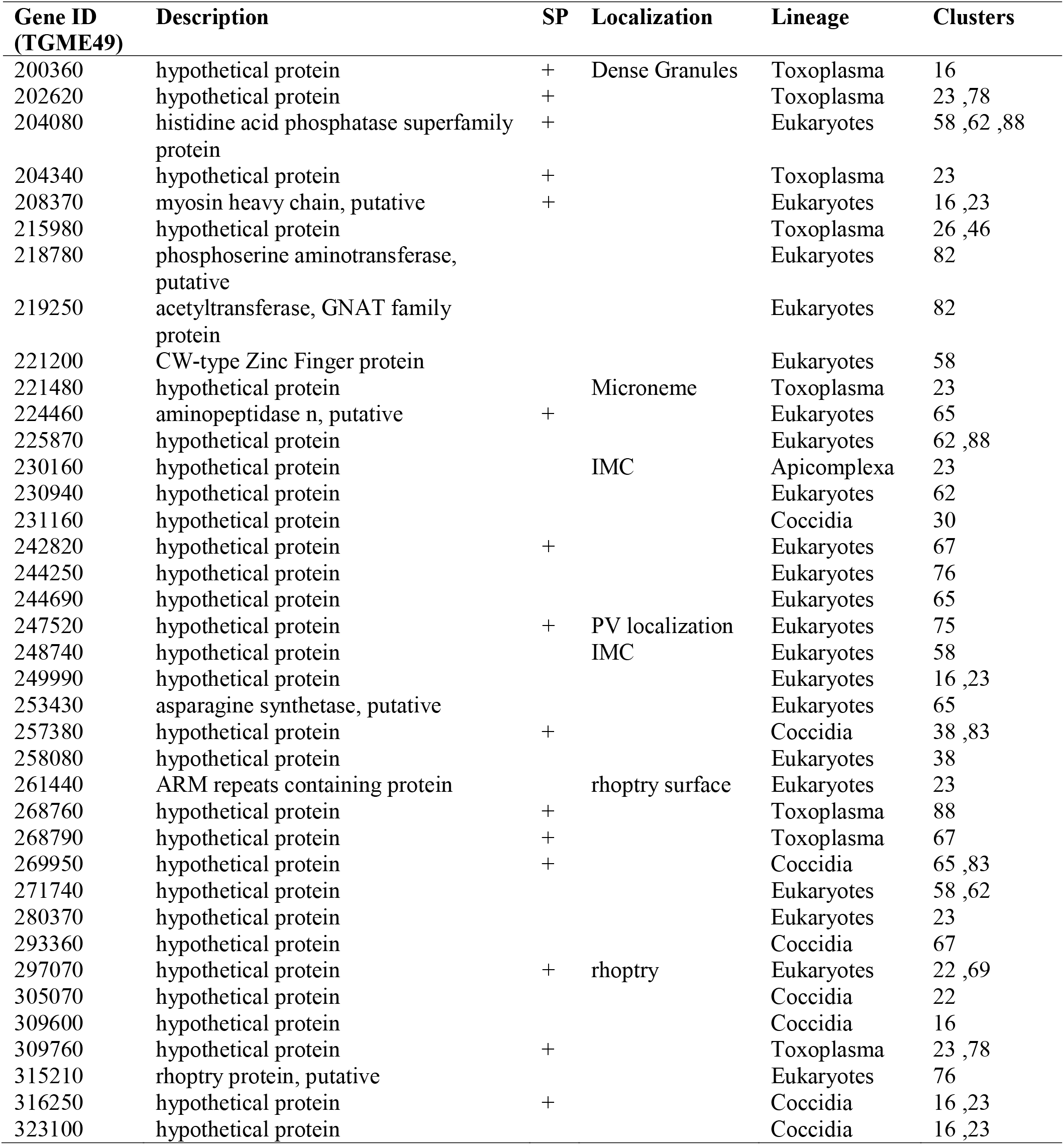
Putative Novel Invasion Factors. Columns indicate Gene IDs (TGME49), their product description, the prediction (+) of a signal peptide (SP), localization information from ToxoDB, predicted lineage and cluster(s) in which it occurs.

Focusing on specific complexes, cluster 16 predicts a novel dense granule complex, with many proteins annotated as localized to the dense granule (e.g. GRA12, GRA32, GRA35, GRA44 and a MAF1 homolog), together with several novel invasion proteins. For example, TGME49_200360 is a hypothetical protein that has been previously localized to the parasitophorous vacuole (PV)^73^. The interacting proteins, GRA35 and GRA44, are also present in the overlapping cluster 78, suggesting that they more represent core subunits of both complexes. Cluster 67 is a dense granule complex containing GRA37, MAG1 and three hypothetical proteins. MAG1 is a known component of the cyst wall during the bradyzoite stage; however, none of the other proteins identified in this cluster were a recent proteomic survey of cyst wall components^74^. The presence of MAG1 in this complex might therefore represent a novel tachyzoite-specific function for this protein. Cluster 75 highlights four proteins that form an isolated subnetwork (i.e. they do not interact with any other network proteins) and include the known rhoptry proteins ROP40, ROP11 and ROP47. This cluster has the highest average SNP score and contains the two network proteins with the highest SNP score, ROP47 and the hypothetical protein TGME49_247520, within the cluster (**Figure 5A**). Additionally, this cluster has the third highest average expression correlation (Pearson correlation = 0.833; **Supplemental Table 7**), suggesting that its function is tightly regulated. Cluster 76 also predicts a novel rhoptry complex that contains two relatively uncharacterized rhoptry proteins, ROP14 and RON1, and two hypothetical proteins. Integration of SNP scores reveals clusters 67, 75 and 78 are among the top ten clusters with the highest average SNP score. The distribution of SNPs for proteins in these clusters across previously designated clades A-F^22^ (and a group of 18 unclassified strains) is visualized within node pie charts (**Figure 5A**). These can be used to generate hypotheses regarding the conservation of these complexes. For example, the high proportion of SNPs present in clade E in the protein TGME49_247520 in cluster 75 suggest that associations with this protein might not be conserved and contribute to variation in the pathogenic phenotype of this relatively small clade of four strains.

**Figure 5.**
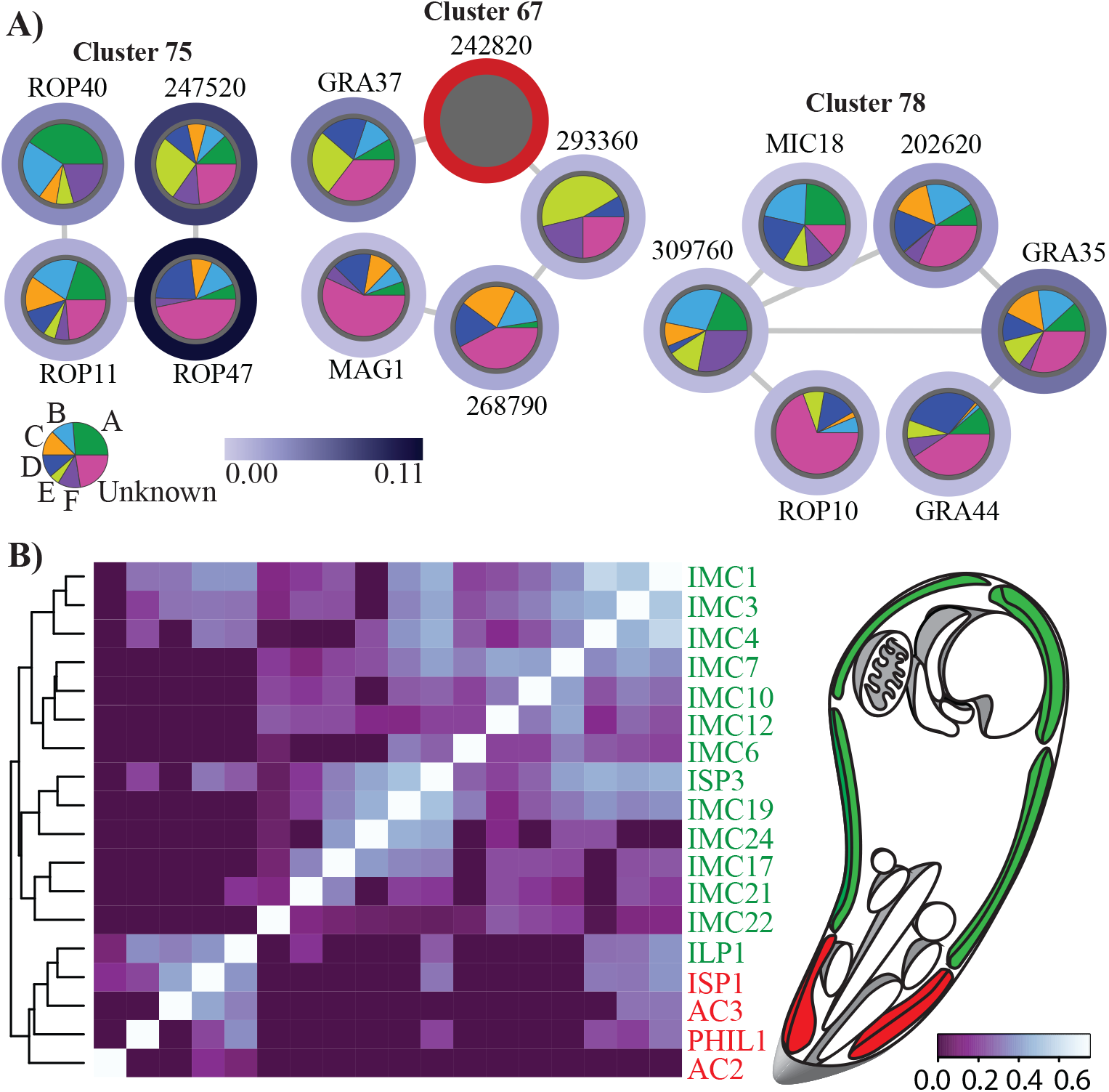
(A) These three putative invasion complexes are among the top ten clusters with the highest average SNP score. The proportion of SNPs present in each clade for each protein is indicated by the pie chart within each node. SNP scores for proteins, such as TGME49_242820, without reliable annotation in the ME49 reference genome were not calculated and are indicated with a red border. The legend indicating clade classification displays the proportion of 80 strains belonging to each of the 7 categories. (B) Heatmap showing co-elution relationships between IMC proteins. For each protein pair, the highest pcc score from each co-elution experiment for IMC proteins was used to generate a matric of co-elution relationships. Clustering of these relationships recapitulates the sub-compartmental structural organization of the IMC. Only proteins with five or more spectral counts are considered. Proteins previously localized to the apical and basal/central subcompartments are highlighted in red and green, respectively. Hierarchal clustering was performed using the complete linkage method and heatmaps were generated using heatmap.2 function in R.

### Putative IMC complexes recapitulate known structural organization and predict novel IMC-related proteins

Clusters 48, 71, 74, 77 and 89 represent putative IMC complexes and predict five novel IMC proteins (**Table 3)**. Two of these proteins have been described as localized to the IMC in ToxoDB user comments, with TGME49_254870 specifically localized to the apical complex. To examine whether the rigid structural hierarchy of distinct apical, central and basal compartments was recapitulated in the co-elution datasets, matrices of the highest pairwise PCCNM or WCC in any experiment were computed for all pair-wise IMC proteins. The highest pair-wise scores were utilized to account for potentially missing data points. Hierarchical clustering of these matrices successfully reconstructs this spatial sub-compartmentalization with proteins from the basal/central and apical compartments segregated into distinct clusters (**Figure 5B; Supplemental Figure 7**). The notable exception is ILP1 which has been previously reported to localize to the central compartment^73^. These results support the network’s ability to predict IMC interactions that are largely consistent with its structural organization. Cluster 48 contains ISP3, IMC19 and IMC22. The spatial proximity of IMC19 and ISP3 has already been confirmed by BioID^61^ while all three IMC proteins have been localized to the central subcompartment of the IMC^61, 75^. Cluster 71 also recapitulates known IMC architecture in that both PHIL1 and ISP1 are localized to the apical subcompartment^75, 76^. Cluster 74 represents a novel complex that contains IMC17, a centrally localized protein, and two uncharacterized proteins: the IMC-localized TGME49_242070^77^ and TGME49_267430. Together these results demonstrate the efficacy of ToxoNet at recapitulating meaningful interactions in the IMC.

**Table 3:** Putative Novel IMC Proteins. Columns indicate Gene IDs (TGME49), their product description, the prediction (+) of a signal peptide (SP), localization information from ToxoDB, predicted lineage and the cluster(s) in which it occurs.

| **Gene ID**<br>**(TGME49)** | **Description** | **SP** | **Localization** | **Lineage** | **Clusters** |
| --- | --- | --- | --- | --- | --- |
| 220390 | hypothetical protein |  |  | Coccidia | 48 |
| 232600 | phospholipase, patatin family protein | + |  | Apicomplexa | 48 |
| 254870 | hypothetical protein |  | Apical<br>Complex | Coccidia | 48 |
| 267430 | DnaJ domain-containing protein |  |  | Eukaryotes | 74 |
| 242070 | cAMP-dependent protein kinase<br>regulatory subunit |  | IMC | Eukaryotes | 74 |

## DISCUSSION

ToxoNet represents the first genome-scale protein-protein interaction network for *Toxoplasma gondii*. Based on co-elution data for 1423 proteins across 6 experiments, we applied two machine learning strategies to integrate additional functional genomics datasets, including gene co-expression, phylogenetic profiles, and domain-domain interactions, to lend support to predicted interactions (**Figure 1**). The first strategy relies on a standard *supervised* approach, in this case Random Forest, which requires gold standard training datasets of known positive and negative interactions. Traditionally such training datasets have been challenging to generate for non-model organisms, typically relying on the inference of orthologous interactions that have been characterized for model organisms. Consequently such datasets tend to be enriched for highly conserved proteins. In an attempt to overcome potential biases that such training data may generate, we therefore adopted a second strategy based on an *unsupervised* approach, in this case Similarity Network Fusion^42^, which instead relies on a non-linear iterative strengthening or weakening of interactions based on their support across multiple complementary datasets. To our knowledge, this is the first instance of applying such an approach to predict protein-protein interactions. While we find that *supervised* approach, in general provides improved performance, we nonetheless find that the *unsupervised* approach performs significantly better that random (**Supplemental Figure 4**). Further, it predicted several previously characterized interactions that were not captured by the *supervised* approach. We therefore suggest that future protein-protein interaction studies also consider applying a similar two-pronged strategy.

The resulting network consists of 2,063 high quality interactions between 652 proteins. While this number falls short of the 8,920 predicted proteins from the *T. gondii* ME49 genome, previous screens of the tachyzoite stage have identified 2,252 proteins expressed at this stage^78^. In this study, the sonication-based lysis conditions were optimized for soluble proteins. Proteomic screens of tachyzoite membrane proteins have identified 841 proteins^79^. The underrepresentation of membrane proteins likely accounts for the 1,423 proteins identified in our fractions. In future studies, this technology can be adapted to optimize membrane complexes with non-ionic detergents or to investigate different lifecycle stages to increase coverage of the *T. gondii* proteome. Comparisons to a recent network of protein interactions generated for the related apicomplexan parasite, *Plasmodium* sp.^41^ shared 22% of the interactions predicted by ToxoNet. This is surprisingly high considering that protein-protein interaction datasets rarely feature high levels of overlap, even in studies within the same organism using comparable methods^80^. Notable apicomplexan complexes predicted in both species, such as the Moving Junction and the Glideosome, are missing from the overlapping set of interactions and are better recapitulated in either the *T. gondii* or *Plasmodium* network, respectively. This suggests that technical differences, such as sample preparation and data processing, still limit cross-species comparisons of protein-protein interaction networks, that might otherwise identify instances of network ‘rewiring’ underlying species-specific adaptations to their specialized life cycles.

Application of the graph clustering algorithm, ClusterONE, predicts 93 clusters that recapitulate known, as well as novel protein complexes. Novel complexes successfully cluster proteins with known functional and spatial relationships, such as the organization of proteins that mediate pathogenesis, or localize to compartments, such as the mitochondria, cytosol and nucleus. It is noteworthy that one-third of the proteins identified by mass spectrometry are indicated as hypothetical by ToxoDB^81^, and their identification in this co-elution study provides evidence that they are expressed in the tachyzoite stage. Many of these hypothetical proteins and other poorly characterized proteins are present in protein complexes, representing valuable opportunities to drive new discoveries. Despite the limitations associated with generating a network on a non-model organism, the *T. gondii* network demonstrates robust congruence with parasite biology as demonstrated by benchmarking of additional datasets. This is evident with high correlation of transcript expression between pair-wise interactions, organization of essential proteins and agreement between spatial proximity of proteins in the cell and in the network.

Detailed analyses of identified protein complexes yield a number of novel insights into the organization of complexes with implications for both specialized parasite processes such as host invasion, as well as parasite-specific adaptions of otherwise highly conserved pathways. For example, many rhoptry and dense granule proteins are known to mediate important roles in maintaining the intracellular tachyzoite lifecycle, co-opting host machinery and regulating the host immune response. However, the full complement of these proteins remains to be elucidated and many that have been identified remain uncharacterized. Among these clusters are known invasion complexes, such as the MIC1/4/6 and GRA2/4/6 complexes, that additionally contain a number of novel components. These data also yield additional insights into the function of previously uncharacterized invasion factors, including GRA32, GRA37, GRA38, ROP11, ROP14, and ROP40, as well as predict many novel invasion proteins. Similarly, ToxoNet also provides additional insights into the function and organization of the alveolate-specific IMC, an important organelle for host cell invasion and sexual reproduction^7^. Our co-elution data reconstructs the rigid sub-cellular organization of this compartment and as such, yields new testable hypotheses concerning the sub-cellular localization of novel IMC-related proteins and their complexes. Beyond these specialized pathways, several recent studies featuring fitness and proteomic screens, have identified an increasing number of apicomplexan-specific proteins that localize to the mitochondria, suggesting that this otherwise highly conserved organelle features a number of parasite-specific adaptations, particularly with respect to ATP synthase and oxidative phosphorylation^82–85^. This expanded view of the apicomplexan mitochondria is reflected in ToxoNet with the prediction of clusters that are supported by these previous studies. Here we identify and localize another novel apicomplexan-specific mitochondrial protein, TGME49_306650, that was previously uncharacterized. Its clustering with known ATP synthase and oxidative phosphorylation machinery suggests a role in these processes.

The construction of protein interaction networks provides a valuable scaffold onto which additional metadata may be integrated. For example, previous studies have leveraged protein interaction networks to inform on properties of essentiality and conservation, domain architectures, as well as taxon-specific representations of microbiome functionalities^35, 86, 87^. Here we show how ToxoNet can be used to interpret genetic variation information derived from 80 strains of *T. gondii*. Such approaches enable strain-level insights into the organization and function of protein complexes. In particular these visualizations allow us to distinguish proteins or protein complexes that represent conserved functions from those that underlie strain-specific functionality, which could be associated with host or tissue tropism.

## CONCLUSIONS

Here we present ToxoNet, the first high quality protein interaction network for *T. gondii*. We validate the quality of the network through systematic comparison of other protein interaction networks, both from other organisms as well as through the application of complementary technology. Our network predicts 93 complexes, including well characterized complexes as well as complexes containing novel components and novel complexes. These data reveal a wealth of testable hypotheses and we therefore make the dataset freely available as a community resource through ToxoDB^88^.

## MATERIALS AND METHODS

### Culturing *T. gondii ME49* parasites and protein extract preparation

*T. gondii* ME49 parasites were cultured in two batches for 3 days in human foreskin fibroblast (HFF) cells supplemented by D10+M199 media. Parasites were harvested, washed with PBS and pelleted at 1500g at 4°C for 15 min. The cells were frozen at -80°C. The pellet was thawed and re-suspended in 1200 µl PBS and 1mL lysis buffer (10mM Hepes-pH 7.9, 1.5mM MgCl_2_, 10mM KCl) with 1mM DTT and Roche mini protease inhibitor cocktail. It was sonicated for a total processing time of 8 min 50 sec with 10 sec on, 10 sec off cycle at 30 - 35 W and centrifuged at 4C and 2000g for 20 min yielding a total supernatant of 760 ul. Protein concentration was determined by absorbance at 280nm to be 31.0 ug/ul.

### Pre-enrichment before HPLC Fractionation by affinity beads

We used affinity beads (NuGel^TM^ *PRO*spector^TM^) to pre-enrich *Toxoplasma gondii* lysate to capture five distinct sub-proteomes. In this experiment, we added one volume of Cleanascite^TM^ *PRO* to five volumes of the sample to remove lipids and any insoluble biomass. We then added PRO-BB binding buffer (pH 6.0) to the delipidated samples in a 1:1 volume ratio. The resulting mixture was then added to different reagent beads (*PRO*-A, *PRO*- B, *PRO*-C, *PRO*-L, *PRO*-R from NuGel^TM^ *PRO*spector^TM^ tool kit) in the Spin-X filterers (from NuGel^TM^ *PRO*spector^TM^ tool kit). The samples and beads were mixed for 10 min, and then centrifuged to collect the filtrate as ‘flow-through’ fractions. We then added PRO-BB binding buffer to wash the sample. We eluted the bound proteins by 200 ul elution buffer (0.2 M Tris, 0.5 M NaCl, pH 9.0). The buffer was exchanged for HPLC loading buffer by Zeba desalt spin column (Thermo) before HPLC fractionation. The resulting elutes from the five beads were kept for later HPLC fractionation.

### HPLC Fractionation

We fractionated *T. gondii* cell lysate and enriched eluates from affinity beads using ion-exchange (IEX) liquid chromatography by an Agilent 1100 HPLC system (Agilent Technologies, ON, Canada) individually. A PolyCATWAX mixed-bed ion exchange column (200 x 4.6 mm id, 12 μ, 1500 A) was used for whole lysate samples, and a 240 min salt gradient (0.15 to 1.5 M NaCl). For enriched eluate samples, a PolyCATWAX mixed-bed ion exchange column (200 x 4.6mm id, 5 μm, 1000A) was employed. Enriched samples were fractionated into 60 fractions by using a 120 min salt gradient (0.15 to 1.5 M NaCl). We collected the fractions every 2 mins. As a result, 120 fractions were collected for the whole cell lysate, and 60 fractions of each eluate from affinity bead. In total, 420 fractions were collected.

### LC-MS/MS analysis

All HPLC fractions were precipitated, re-dissolved and then digested by trypsin overnight at 37 °C resulting in peptides, which were subsequently dried and re-dissolved into 5% formic acid before LC-MS/MS. The LC-MS/MS was performed by a nano-flow HPLC System (EASY-nLC, Proxeon, Odense, Denmark) coupled with a LTQ Orbitrap Velos Mass Spectrometer (Thermo Fisher). First, the peptides were loaded into a 2.5 cm trap column (75 mm inner diameter), which was packed with Luna 5u C18, 100A beads (Phenomenex), by an auto-sampler. Next, a 10 cm analytical column (75 mm inner diameter), packed with 2 mm Zorbax 80XDB C18 reverse phase beads (Agilent), was connected to the trap column for peptides separation. A 60 min gradient of CAN in water (1% formic acid) from 5% to 35% was used to elute peptides. Electro-spray ionization was set at 2.5kV, and the mass spectrometer was operated in a data dependent mode (One full MS1 scan followed by MS2 acquisitions on top 10 precursor ions). The fragmentation was performed by 35% normalized collision energy at CID mode.

### Protein Identification and Label Free Quantification

We converted all raw files generated from LTQ Orbitrap Velos Mass Spectrometer to mzXML files by ReAdw software. The FASTA file was downloaded from ToxoDB v11^88^ and common contaminant peptides and corresponding reverse sequence decoys were added for false-discovery rate (FDR) evaluation. We used SEQEST algorithm as protein searching engine and set the FDR less than 1% for both peptide spectral matching and protein matching for each sample. We then used STATQUEST model to probabilistically evaluate the confidence score to all matching. All the fractions are then compared and concatenated by CONTRAST software tool.

### Capturing similarity of coelution profiles

The similarity of coelution profiles for two proteins was estimated using various metrics. Three metrics implemented by Havugimana et.al were used as specified in ^37^: a) pearson correlation coefficient with noise modeling (PCCNM) – which introduces random noise into the profiles in order to negate the high spurious correlations arising due to low abundance; b) weighted cross correlation (WCC) – which negates minor spectral shifts arising in the coelution profiles during collection of fractions; c) overall coapex score (Coapex1) – which captures the number of co-elution experiments in which the same fraction contains peak abundance for both proteins. In addition, three other scores were implemented: a) mutual information (MI), implemented using the “entropy” package in R, in order to capture non-linear dependencies between the two coelution profiles; b) topological overlap similarity (TOM) – typically used in gene co-expression networks ^89^ to measure the relative interconnectedness between two proteins by combining the similarities between the two proteins in light of the (captured via their PCCNM, WCC scores) along with those of its shared neighbours – implemented using the scran and WGCNA packages in R; and c) individual coapex score (Coapex_X) – capturing the number of fractions with same peak abundance for both proteins in an experiment. In summary, for each protein pair, 6 PCCNM scores, 6 WCC scores, 6 Coapex_X scores, 6 MI scores, 6 TOM-PCCNM, and 6 TOM-WCC scores are generated, corresponding to each of the 6 experiments (five from bead purifications and one from whole cell lysate). Further, apart from Coapex1 (which is calculated over all experiments), overall scores were calculated using two approaches for the other measures by capturing similarity over all experiments (420 fractions) after ensuring that the two proteins coelute in at least one experiment: a) Calculate similarities over 420 fractions (PCCNM_overall, WCC_overall, MI_overall, TOM-PCCNM_overall, TOM- WCC_overall, MI-overall); and b) Calculate similarities only over the experiments in which either / both of the proteins are detected (PCCNM_overall-E, WCC_overall-E, MI_overall- E). Note that the latter was not carried out for TOM-based scores which considers shared neighbours in each experiment, resulting in too many combinations of experiments, with several being populated sparsely. Details of scores are shown in **Supplemental Table 3**.

### Integrating functional genomics datasets

The coelution datasets are integrated with additional functional genomics data for *T. gondii* using a supervised machine learning classifier in order to generate a protein interaction network. The additional datasets are:

(a) Domain-domain interactions: Lee et. al.^46^ generated log-likelihood scores to capture how often two Pfam domains^90^ found in two proteins are involved in physical interaction. An integrated log-likelihood score was generated for each protein pair based on the log-likelihood scores assigned for their Pfam domain.
(b) Phylogenetic profiles – Phylogenetic profiles represent the pattern of gene/protein distribution in sequenced genomes. Phylogenetic profiles for *T. gondii* were obtained from Phylopro v2^91^, which stores gene/protein distribution for 165 eukaryotic proteins^92^. Two types of phylogenetic profiles were generated: i) where presence/absence of orthologs is represented as 1s and 0s ii) where presence of homolog is represented in terms of modified BLAST E-value reflecting extent of sequence similarity, thereby capturing more information than binary gene presence/absence^93^. The similarity of phylogenetic profiles of two proteins was estimated using mutual information, implemented by the entropy package in R.
(c) Co-expression – Co-expression of *T. gondii* proteins in 44 conditions was obtained from 8 co-expression datasets, from ToxoDB v11^81^. The datasets were split into two based on the method: RNAseq and Microarray. The similarity between two coexpression profiles was estimated using pearson correlation coefficient. The datasets considered are: Buchholz_Boothroyd_M4_in_vivo_bradyzoite_rnaSeq, Sibley_ME49_bradyzoite_rnaSeq, DBP_Hehl-Grigg_rnaSeq, Knoll_Laura_Pittman_rnaSeq, microarrayExpression_Boothroyd_LifeCycle, microarrayExpression_Matrajt_GSE23174_Bz, microarrayExpression_Roos-Tz, microarrayExpression_Sullivan_GSE22100_GCN5-A.
(d) Gene fusion and Textmining – Protein pairs in *T. gondii* reported to be functionally interacting according to gene fusion and text-mining (co-citation) evidence from STRING v10 were considered^94^. The scores representing their similarity in the STRING database were considered as is.

### Supervised machine learning approach to network generation

A supervised machine learning approach based on a RandomForest classifier (implemented using weka software suite v3.6) was used to integrate the similarity scores of coelution and additional datasets using a training dataset of 511 positive interactions and 1533 negative interactions generated from ToxoCyc^95^ and 1:1 orthologues of manually curated complexes from Cyc2008 (http://wodaklab.org/cyc2008/), CORUM^96^ (**Supplemental Table 4**). The positive training dataset consists of 333 pairs from Toxocyc, 133 pairs sharing same GO-Biological Process terms (GO term annotation by Pfam2GO taken from^22^), 206 pairs from orthologs of yeast complexes in Cyc2008, and 516 pairs from orthologs of human complexes in CORUMCore (complexes with 50 members). Duplicate pairs and ribosomal pairs were removed from these datasets, yielding an overall positive training dataset of 511 pairs. The negative training dataset was generated by choosing pairs that do not share common localization (according to Apiloc v3 annotation wherever available (152 pairs), or by comparison of GO terms of orthologs in Cyc2008 and CORUM otherwise (2227 pairs and 1912 pairs respectively)). Of the 61 features in total comprising of both co-elution and functional dataset similarity scores, feature selection identifies 33 attributes as providing information gain with respect to the training data (**Supplemental Table 3**). Based on 10-fold cross validation, the RandomForest classifier is able to classify the training datasets with an AUC score of 0.811 (**Figure 1**). This RandomForest classifier was used to evaluate the set of 174455 protein pairs with biochemical support (co-fractionation score ≥ 0.5 in any of the experiments). The resulting set of predicted interacting pairs were further pruned to: i) remove pairs containing proteins with a spectral count of ≥ 5 (to remove spurious correlations arising out of low abundance - 90% of the scores with a value of –nan according to PCCNM have <=5 spectral counts in one of the protein pairs); and ii) remove pairs of proteins not containing any unique peptides distinguishing one from the other.

### Unsupervised machine learning approach to network generation

An unsupervised machine learning approach, based on Similarity Network Fusion (SNF)^42^, was also adopted for integrating the similarity scores of coelution and additional datasets. For a set of 1329 proteins common to the cofractionation, coexpression and phylogenetic profile datasets, seven similarity networks were generated based on three similarity measures for coelution data (PCCNM_overall, WCC_overall, Coapex1), two similarity measure for coexpression data (COEXPR_RS, COEXPR_MA), and two similarity measures for phylogenetic profiles (MI based on presence/absence (MI-PresAbs) and sequence similarity metric (MI-pij)). Different combinations of the seven similarity networks were integrated using SNF for a range of parameters (K – 2 to 30, alpha – 0.2 to 1, I – 2 to 30): all scores, removing COEXPR_MA, removing MI-PresAbs, removing MI-pij. The resulting set of integrated pairs were pruned at a false positive rate of 0.05 (determined based on the negative training dataset generated for the supervised learning technique). These pairs were further pruned in a manner similar to the supervised network in order to i) remove pairs containing proteins with a spectral count of ≥ (to remove spurious correlations arising out of low abundance) ii) remove pairs where there are no unique peptides distinguishing one protein from the other and iii) remove pairs with a co-fractionation score ≤ 0.5 in all experiments. The predicted interaction sets from each of the permutations were clustered using MCL (at an iteration of 2.6). The congruence of the predicted networks in identifying biologically relevant clusters was evaluated by estimating the similarity of the clusters with a reference set of complexes in the training dataset (pertaining to orthologs of CORUM/Cyc2008 complexes and members of same pathway in ToxoCyc). The similarity was estimated using the Bader-Hogue score^66^ and all clusters with a cutoff ≥0.25^65^ were considered to be similar. Finally, all protein pairs with a PCC ≥ 0.5 in COEXPR_RS were considered in order to generate the high confidence unsupervised network (**Supplemental Table 6**).

### Generating combined network and high-confidence network

The Bader-Hogue similarity score^66^ was used to identify the best unsupervised predicted networks in terms of their ability to identify biologically relevant clusters: From the various unsupervised predicted networks generated for different combinations of similarity matrices and parameters, the network with the highest number of clusters according to the Bader- Hogue cutoff score (≥ 0.25) was selected. For the supervised network, the network generated based on feature selection was chosen. These two networks were integrated to generate the overall combined network. Further, to generate a high confidence set, the supervised network was derived by considering interactions greater than a score corresponding to a false positive rate of ∼0 (indicated by a * on the ROC curve – Figure 1). For the unsupervised network, the high confidence version was generated by considering all those interactions that are ≥0.5 for COEXPR_RS). These two high confidence networks were collated in order to generate a high-confidence network.

### Network analysis

Topological analysis of the network was achieved using the Cytoscape plugin NetworkAnalyzer to determine node degree, node betweenness, network degree distribution, network characteristic pathlength and network shortest pathlength distribution. Essentiality information was integrated from a genome-wide CRISPR screen of the tachyozoite lifecycle^82^. Essential proteins were defined by a phenotype score of less than or equal to -2 based on the reported separation of proteins with experimentally validated essential and dispensable fitness phenotypes^24^. The *T. gondii* network was compared to previously published *T. bruceii*^62^ and *Plasmodium*^41^ networks using 1:1 orthologs identified by InParanoid v4.1^97^. Information from spatial proximity experiments was analyzed in the context of the network. In publications that utilized both BioID and APEX techniques^58, 59^, the subnetwork was defined by the network proteins present in the overlapping set detected by both methods. In cases where only BirA* fusion proteins were utilized, the subnetwork was defined by network proteins detected in streptavidin pull-downs that were not present in negative controls^60, 61^. The Python package networkx was used to calculate the pairwise pathlength of all subnetwork proteins and the average was the characteristic pathlength of the subnetwork. The 55 network proteins that are isolated in clusters of 2-4 proteins were not included in this analysis due to their disconnection from the main network. Subnetwork permutations were generated by randomly selecting the same number of proteins from the network and determining the characteristic path length. This was repeated 1000 times for each subnetwork. P-values were determined using the pnorm function in R. For the G.O. comparisons, computed G.O. component, processes and function terms were taken from ToxoDB. Of the number of interactions where both proteins were annotated, the percent of interactions with the same term was calculated. Network permutations were generated by randomizing the target and source lists. For each 1000 random networks, the percent of interactions with the same term was calculated. The significance of the difference between the network value and the distribution of random values was determined using the pnorm function in R. To analyze coexpression correlation of network interactions, RNAseq data was taken from ToxoDB from the following directories: tgonME49_Boothroyd_oocyst_rnaSeq_RSRC, tgonME49_Gregory_ME49_mRNA_rnaSeq_RSRC, tgonME49_Reid_tachy_rnaSeq_RSRC. The total 11 timepoints were merged from the profiles.min files and the Pearson correlation values were determined for network interactions, all combinations of network proteins that were not predicted to be interacting, and 10,000 random combinations from the full proteome. The significance of the difference in the distributions of these respective values was determined by a two-sample Kolmogorov-Smirnov test.

### Clustering and Cluster Analysis

Clusters were identified in the high confidence network using ClusterONE^65^ using a density parameter of 0.25. The density parameter was optimized by iteratively scanning each possible value. To select a final cluster set, the number of recapitulated complexes utilized in supervised training data with an overlap score of 0.25 or greater^66^ was calculated. The average number of recapitulated training complexes was also calculated from 100 random cluster permutations. The set of clusters that recapitulated the greatest number of training complexes minus the average number of training complexes recapitulated at random was selected as the final set of clusters. For the analysis of coexpression of cluster proteins, the average cluster coexpression correlation was calculated for each cluster by averaging the Pearson correlation for each pairwise combination of cluster proteins. The average correlation with network proteins was calculated by averaging the Pearson correlation for each pairwise combination of cluster proteins with all network proteins. The p-value was calculated using a Welch’s T-test to determine the significance of the difference of the average coexpression correlation of cluster proteins with each other relative to all network proteins. For the analysis of distribution of essential proteins in clusters, the fraction of essential proteins for each cluster was calculated. 100 random cluster permutations were generated as previously described. The significance of the difference in the distributions of essential proteins in clusters and random clusters was determined by a two-sample Kolmogorov-Smirnov test.

### SNP Analysis

The dataset of SNPs used here compiles sequencing results from previous studies of *T. gondii* genetic diversity^22, 69^, and contains 1,054,454 SNPs spanning 80 different strains of *T. gondii*.

The effect of these substitutions on the ME49 reference genome were assessed using SnpEff v4.3^98^, which produced multiple functional annotations (including the affected gene and type of mutation) for each given SNP. SNPs with minor alleles only present in 3 or fewer strains were excluded from further analyses. The SNP dataset was further filtered down by considering only those SNPs causing non-synonymous mutations for proteins in the network, resulting in a final set of 2893 SNPs. To determine which of these SNPs have pronounced biological effects, the 2881 network SNPs annotated as missense mutations were each assigned a score using PROVEAN v1.1.3. SNPs with a PROVEAN score of -2.5 or below were considered to be putatively deleterious. The SNP score was calculated for each network protein as the number of non-synonymous SNPs normalized by the protein length. The deleterious SNP was similarly calculated, but including only non-synonymous SNPs that were flagged as deleterious by PROVEAN. The previously described neighbor network^22^ designates 62 of these *T. gondii* strains in clades A-F; the remaining 18 are designated as unknown. For each of the non-synonymous SNPs, of the strains carrying the minor allele, the proportion belonging to each clade was calculated. The proportions for each clade were averaged for all these SNPs within a protein to determine the final clade distribution visualized within node pie charts. The average SNP score of each cluster was determined by averaging the SNP score of each cluster protein.

### IMC Reconstruction

To relate cofractionation datasets to characterized structural hierarchies of the apicomplexan IMC, PCCNM or WCC matrices were computed for all known IMC proteins contained in the proteomic data with five or more spectral counts. These matrices contained the highest pairwise PCCNM or WCC score from individual fractionation experiments (i.e., five affinity beads and IEX) or from across all 420 fractions, respectively. Hierarchical clustering of these matrices was performed using the complete linkage method and heatmaps and representative dendograms were generated using the heatmap.2 function in R.

### Endogenous Epitope Tagging and IFA

*T. gondii* PruΔ 88Δ gprt strains were used to generate the endogenous 3×HA-tagged strains used for IFA assays. The 3×HA sequence was inserted into the endogenous locus of TGME49_306650 by double crossover homologous recombination using CRISPR/Cas-based genome editing and homology arms of 30 bp to facilitate the 3×HA integration. For all transfections, 6 μ g of a repair oligo (3xHA).

Parasites were transfected and cloned by limiting dilution after the first lysis. Insertion of 3xHA tag was confirmed by conventional PCR and IFA. All strains were maintained in HFF cells grown in Dulbecco’s modified Eagle’s medium (DMEM) supplemented with 10% heat-inactivated fetal bovine serum (FBS), 0.25 mM gentamicin, 10 U/mL penicillin, and 10 μg/mL streptomycin (Gibco, Thermo Fisher Scientific Inc., Grand Island, NY). For IFA, HFF monolayers were infected with *T. gondii* for 24 hours. Coverslips were then fixed with 0.1% Triton X-100 and incubated with primary antibodies. The signal from primary antibodies was then amplified using species-specific secondary antibodies conjugated to Alexa488, Alexa647 and Rhodamine Red X. DNA was stained using DAPI. Imaging was performed utilizing an Olympus IX81 quorum spinning disk confocal microscope with a Hamamatsu C9100-13 EM-CCD camera (the Imaging Facility at the Hospital for Sick Children).

## ACKNOWLEDGEMENTS

This research was supported by a grant to JP, AE and JB from the National Institutes of Health (R21AI126110), to JP, AE and MG from the Canadian Institutes for Health Research (PJT 152921), and in part by the Intramural Research Program of the National Institute of Allergy and Infectious Diseases (NIAID) at the National Institutes of Health. GS was supported by a student Restracomp fellowship given by the Hospital for Sick Children. Computing resources were provided by the SciNet HPC Consortium. SciNet is funded by: the Canada Foundation for Innovation under the auspices of Compute Canada; the Government of Ontario; Ontario Research Fund - Research Excellence; and the University of Toronto. We would like to thank Peter Bradley (UCLA) for generously providing us with α-F1B ATPase antibody.

## SUPPLEMENTAL INFORMATION

**Supplemental Figure 1:** Precision-Recall curve for the Randomforest approach, for different combinations of features.

**Supplemental Figure 2:** Predicted high confidence supervised network depicted using cytoscape, where proteins are shown as nodes (circles) and the predicted interactions between them are shown as edges (lines). Hypothetical proteins are indicated as squares enclosed by black borders. Well known protein complexes are colored uniquely and encircled – such as ribosome (yellow), proteasome (orange), snRNP complex (light green), glycolytic complex (blue), prefoldin complex (black). Proteins known to be involved in invasion are colored pink, and proteins associated with the IMC are colored purple.

**Supplemental Figure 3**: Fraction of interactions lost for various cutoffs of coexpression measure (PCC) for both the supervised and unsupervised networks.

**Supplemental Figure 4:** Comparison of supervised and unsupervised networks. (A). Venn diagram showing the number of overlapping and unique proteins for the supervised and unsupervised networks. (B) Features of proteins uniquely identified by the unsupervised network: Distribution of spectral counts for the proteins, Box plot comparing the PCCNM scores for the interactions of these proteins with an equivalent set of randomly generated interactions C). Distribution of various cofractionation, coexpression, and phylogenetic scores for Supervised, Unsupervised, and equivalent sets of Random interactions.

**Supplemental Figure 5.** (A) The distribution of random permutations (n=1000) of characteristic path lengths relative to the actual characteristic path length (red line) of network proteins identified in biotinylation BirA-based BioID experiments with GRA13 (n=35, p=0.43), GRA17 (n=48, p=0.1), GRA25 (n=71, p=0.07), ISP3 (n=46, p=0.85), AC2 (n=14, p=0.21). (B) The intersection of interactions predicted in recent *Trypanosome bruceii*^6^ and *Plasmodium falciparum*^7^ protein interaction networks.

**Supplemental Figure 6.** Characteristics of essential and dispensable proteins. (a) Overlapping histograms compare the distribution of the fraction of conserved eukaryotic proteins in essential and dispensable clusters. Boxplots illustrate the fraction of invasion and IMC proteins (b) and hypothetical proteins (c) in essential and dispensable clusters.

**Supplemental Figure 7.** (A) Boxplots compare the distribution of average SNP scores for complexes from different compartments. Invasion complexes include complexes from dense granules, rhoptry and microneme organelles. (B) Hierarchal clustering of the best pairwise wcc score from each coleution experiment for IMC proteins recapitulates the subcompartmental structural organization of the IMC. Proteins previously localized to the apical and basal/central subcompartments are highlighted in red and green, respectively.

**Supplemental Table 1:** Details of peptide counts obtained from mass spectrometry analysis of T. gondii tachyzoites from six co-fractionation experiments

**Supplemental Table 2:** Details of *T. gondii ME49* proteins identified in the cofractionation experiments, along with their elution profiles

**Supplemental Table 3**: Description of scores evaluated in the study along with selected scores from feature selection

**Supplemental Table 4:** Description of POSITIVE and NEGATIVE training datasets used for training the supervised classifier.

**Supplementary Table 5:** Co-elution, coexpression, and phylogenetic profiles of predicted interactions for the supervised network

**Supplemental Table 6:** Co-elution, coexpression, and phylogenetic profiles of predicted interactions for the unsupervised network

**Supplementary Table 7:** Details of Protein Complexes including, protein components, complex size, name of recapitulated training complex, overlap score of training complex, fraction of eukaryotic, apicomplexan, coccidian and toxoplasma-specific proteins, fraction of essential and dispensable proteins, putative localization, average SNP score, average co-expression correlation of cluster proteins, the average co-expression correlation of cluster proteins with all network proteins and the p-value determining the significance complex co-expression.

## Notes

### Competing Interest Statement

The authors have declared no competing interest.

