## Supplementary figures and images for "ToxoNet: A high confidence map of protein-protein interactions in *Toxoplasma gondii* reveals novel virulence factors implicated in host cell invasion"

### Supplemental Figure 1

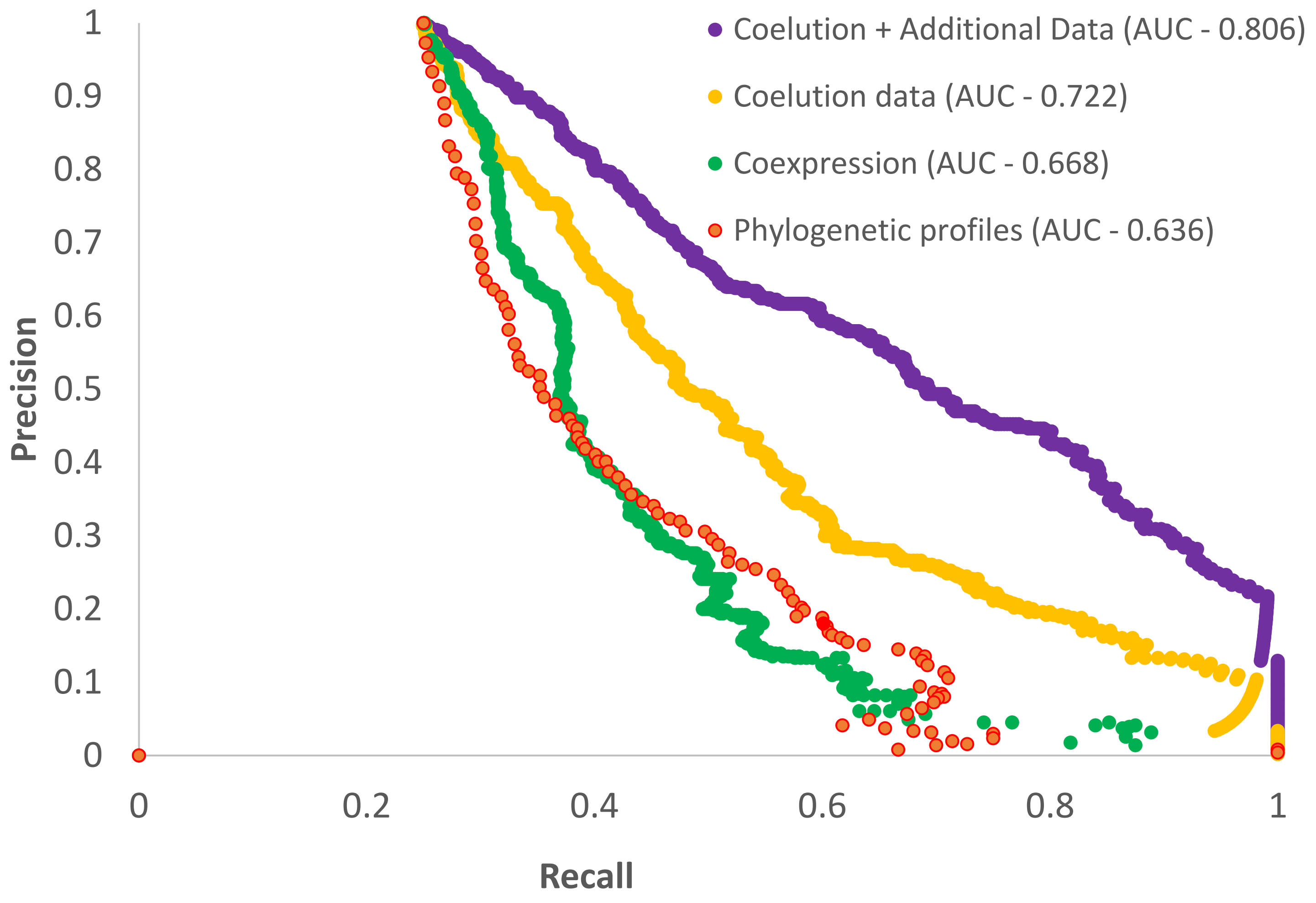

### Supplemental Figure 2

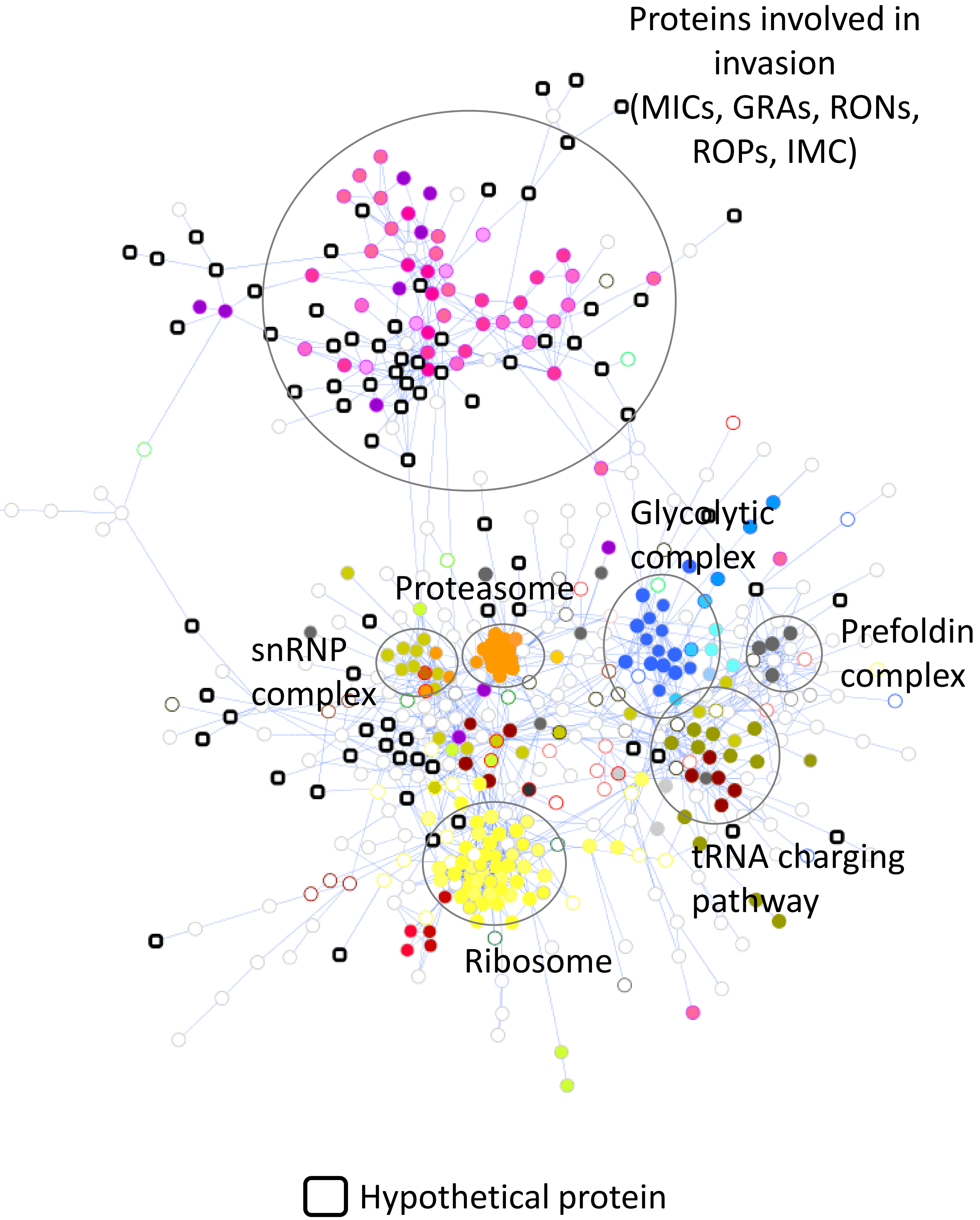

### Supplemental Figure 5

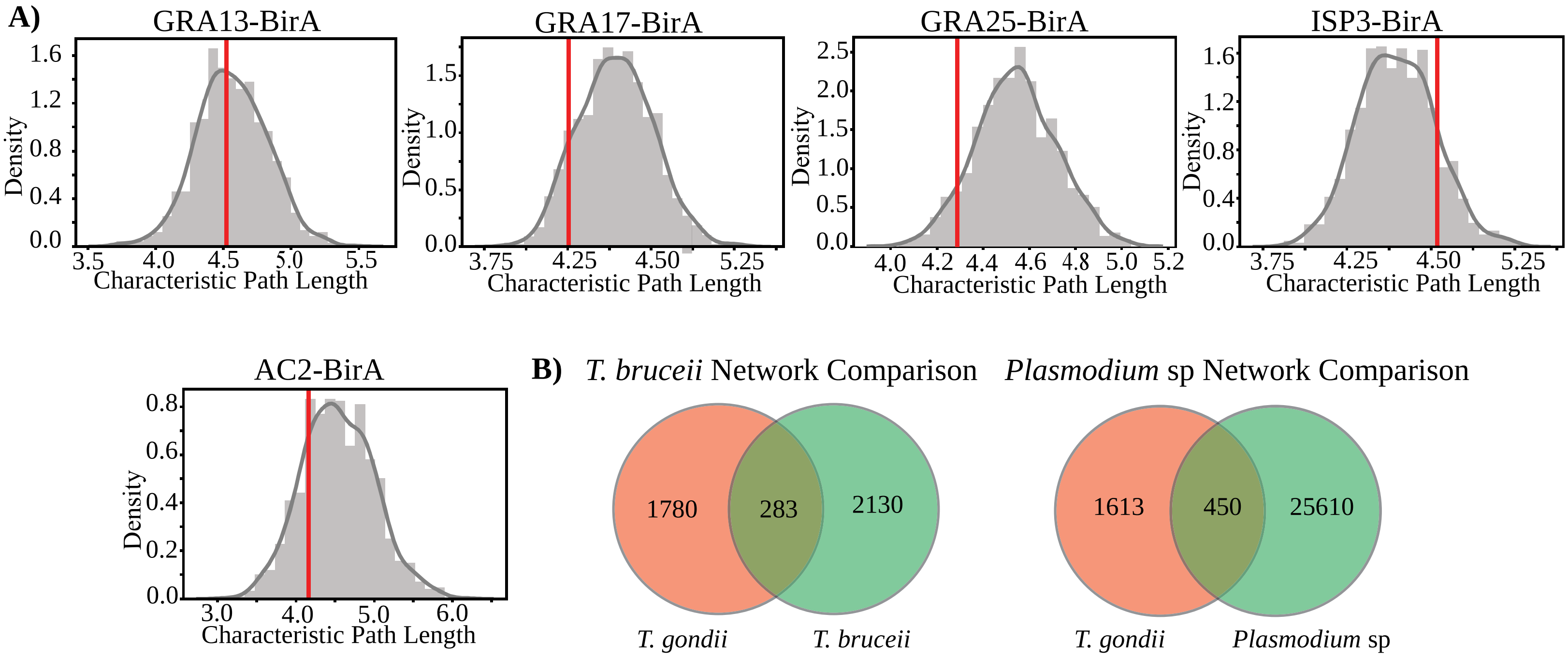

### Supplemental Figure 6

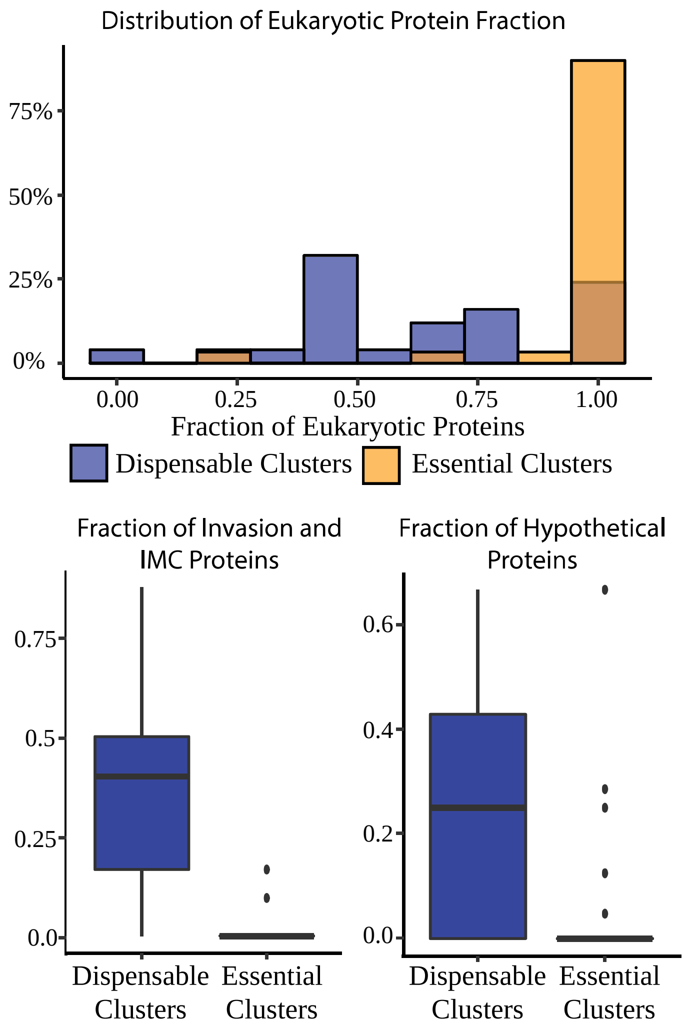

### Supplemental Figure 7

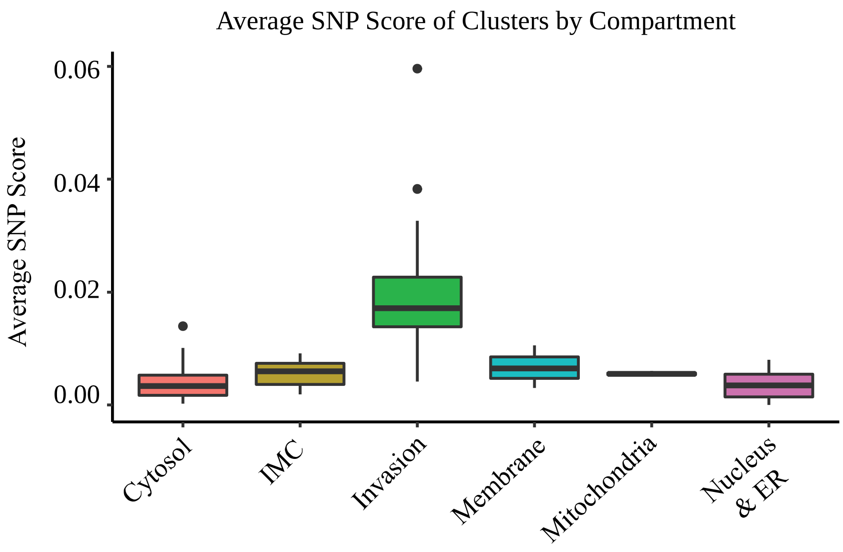

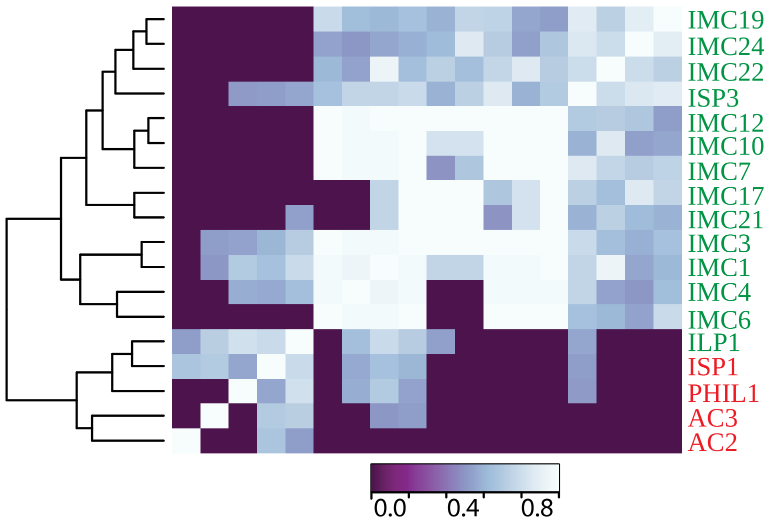
