## Supplemental Figure 3 for "ToxoNet: A high confidence map of protein-protein interactions in *Toxoplasma gondii* reveals novel virulence factors implicated in host cell invasion"

Fraction of interactions lost

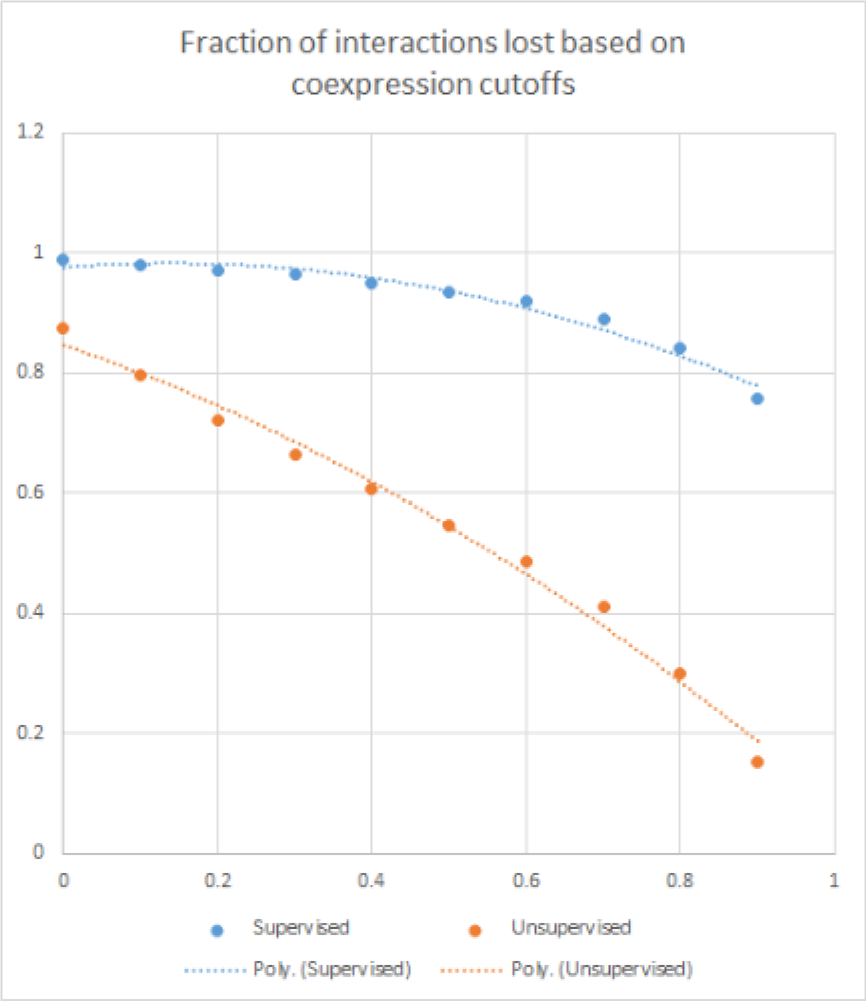


Coexpression cutoffs (based on pearson correlation coefficient)
