## Supplemental Figure 4 for "ToxoNet: A high confidence map of protein-protein interactions in *Toxoplasma gondii* reveals novel virulence factors implicated in host cell invasion"

### A) Distribution of proteins in low confidence network

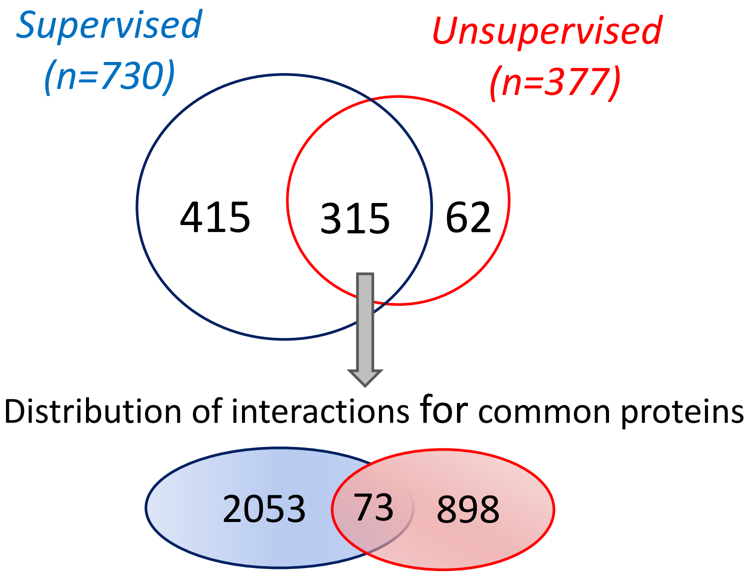


### B) Distribution of spectral counts

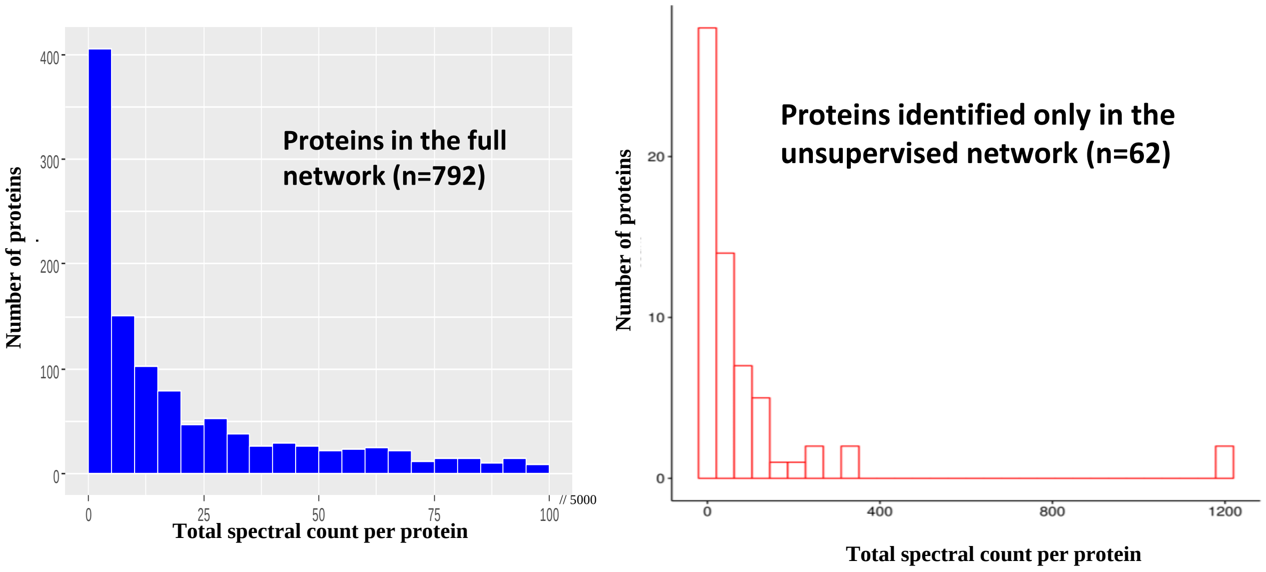


### C) Distribution of scores for Supervised vs. Unsupervised vs. Random Interactions

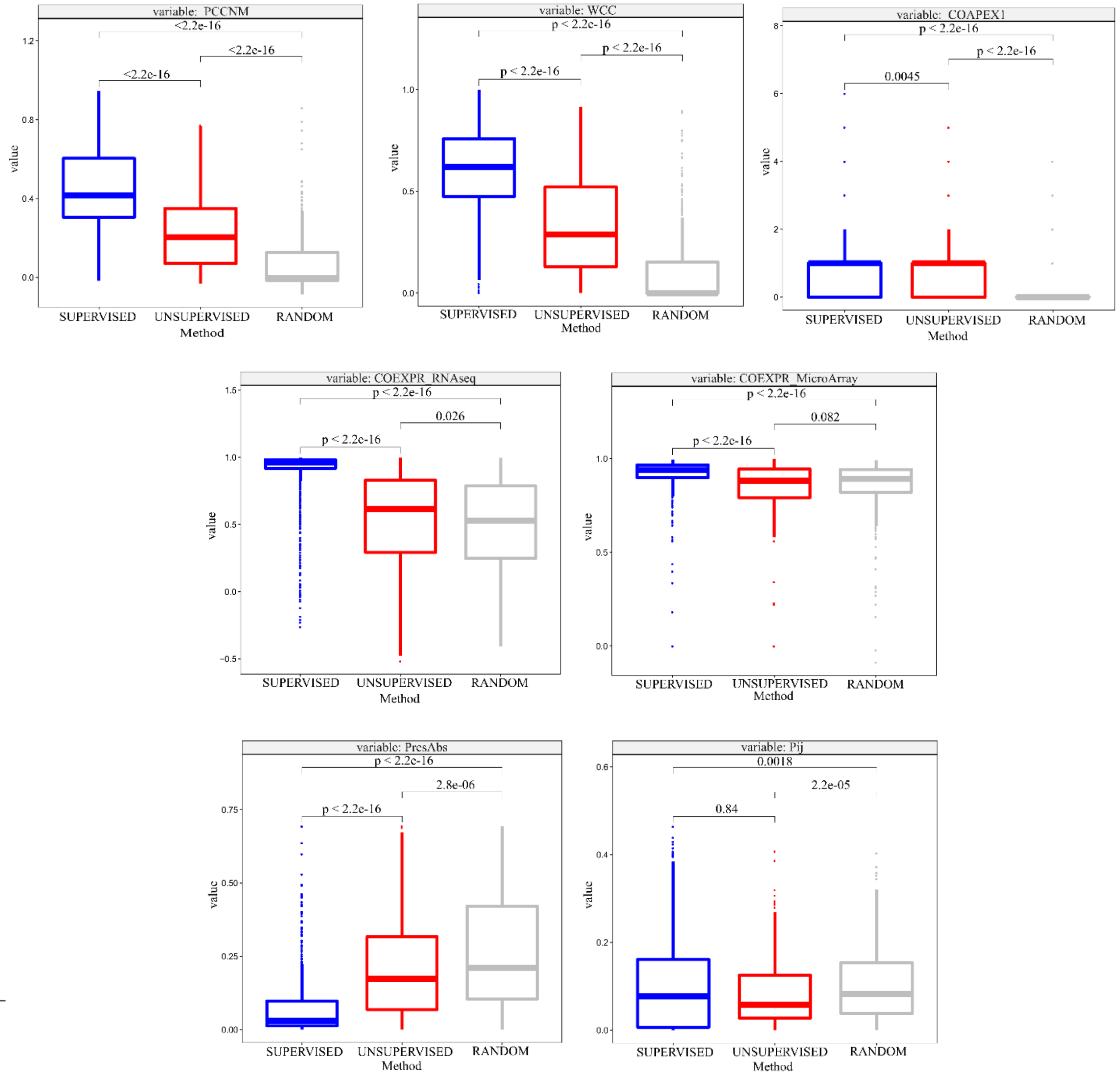
